# Anticipatory emotions and academic performance: The role of boredom in a preservice teachers’ lab experience

**DOI:** 10.1101/2024.01.16.575518

**Authors:** Jesús A. Gómez-Ochoa de Alda, José María Marcos-Merino, Cristina Valares-Masa, María Rocío Esteban-Gallego

## Abstract

Affective experiences within academic contexts significantly influence educational outcomes. Despite this, the literature reveals a gap in generalising these effects to specific classroom activities, partly arising from the absence of suitable instruments to measure emotions in situational educational scenarios. Our study introduces an experience sampling method to measure sixteen discrete emotional states, deriving two scales for positive and negative activating emotions. Grounded in psychological and neuroscientific theories that integrate emotion with cognition, our research explores the interplay between prior knowledge, preservice teachers’ anticipatory situational emotions, and subsequent learning in an experimental science education context. Analysing data from 269 preservice teachers from diverse backgrounds (STEM and non-STEM) at the upper-secondary level, we found that negative activating emotions are often rooted in non-STEM backgrounds and exacerbated by limited prior science knowledge. These negative emotions impact achievement and learning primarily through the mediating role of boredom. Furthermore, our results indicate that the detrimental impact of boredom on achievement is significantly influenced by prior knowledge, with a more pronounced effect on students with lower levels. Given that emotions are amenable to intervention, our findings propose that specifically addressing boredom in students with low prior knowledge could amplify the benefits of educational strategies.

## 1. Introduction

Education constitutes an emotive undertaking. Both students and educators experience emotions triggered by diverse antecedents such as subject matter, cognitive assimilation, pedagogical approaches, and retrospective and prospective achievements. These academic emotions can modulate teachers’ and students’ behaviour [1], influencing their decisions in teaching and learning processes [2], such as time management or learning strategy choice [3]. The emotional states within the classroom significantly influence educational outcomes [4,5]. The Control-Value Theory of achievement emotions (a type of academic emotion related to achievement) posits that the perceived control over achievement activities and the perceived value of these activities and outcomes are the antecedents of achievement emotions, being these emotions the immediate antecedents of learning outcomes [1]. Given emotions’ impact on education, it is advised that teachers consider the affective elements of their pedagogy, particularly in science education, where disengagement often begins in late primary education through to secondary education levels, marked by an increase of negative emotions and a decrease in science valuation and self-efficacy [2]. It is crucial to pinpoint, assess, and understand the sources of student emotions to tailor effective regulatory strategies and foster a conducive emotional climate for science engagement. Furthermore, preservice teachers need to understand their affective influence, as positive emotions towards science can inspire similar sentiments in students, given the transferability of teacher emotions [6].

This study examines the association between preservice teachers’ anticipated emotions before engaging in a laboratory activity and the resulting learning outcomes. Furthermore, we investigate how these emotions may be influenced by their prior educational experiences and knowledge. To initiate the study, we first introduce prevalent misconceptions related to emotions. Subsequently, we present a conceptual framework rooted in neurological and psychological perspectives to understand emotions better, their classification and their effect on learning while emphasising the research gaps that motivated this study.

### 1.1 Preventing common misconceptions

Emotions represent a complex aspect of human behaviour studied across various academic disciplines—psychology, neuroscience, philosophy, anthropology, and sociology—each offering unique interpretative frameworks, methodological approaches, and terminologies. The heterogeneity of these interdisciplinary analyses presents obstacles to the systematic study, research, and communication of emotional phenomena. Notably, the interchangeable use of emotion —physiological response— by feelings —subjective experiences— persists [7] alongside a misapprehension of cognitive and emotional processes as separated mental faculties [8]. This dichotomy, which confronts cognition (traditionally associated with masculinity) in constant contention with emotion (associated with femininity), continues to influence the conceptualisation of human behaviour, thereby perpetuating gender stereotypes within the socio-cultural milieu despite its questionable empirical substantiation [9,10]. The enduring nature of these stereotypes accounts for the neglect of emotion studies in educational settings compared to cognitive studies.

### 1.2 Understanding neurological and psychological conceptual frameworks

From a neurobiological perspective, the Theory of Constructed Emotions [9] posits that "emotions are constructions of the world, not reactions to it". Accordingly, the brain continuously constructs concepts and creates categories to interpret internal and external sensory inputs such as physiological, cognitive, social, and cultural processes. It infers causal explanations for these inputs and predicts action plans in response [9]. Emotions serve as adaptive responses, predicting the most fitting rejoinder based on previous encounters with analogous stimuli, being considered an evolutionary predictive process humans use to subjectively evaluate their environment to estimate the future desirability of various stimuli and respond appropriately [9,11]. When the brain’s internal model constructs a concept related to emotion, this categorisation process culminates in the experience of a specific emotional state. This state is not just a passive reaction but an active construction, integrating both external stimuli and internal cognitive processes. Students actively construct emotions across various tasks, subjects, settings and throughout their academic tenure.

From a psychological perspective, the Appraisal Theory of Emotion [12], in line with the Control-Value Theory [13], posits that emotional states or their constituents arise upon a subjective valuation of stimuli according to their congruence or incongruence with the individual’s objectives and anticipations, the degree of their controllability, and the locus of their origination—whether it be external, internal, or attributable to impersonal factors. In the Theory of Constructed Emotions, these evaluative processes are seen as components of mental representations that anticipate potential actions to situational contingencies. These representations trigger emotional responses (physiological, cognitive, and motor) according to a perception-valuation-action (P-V-A) sequence that can be verbalised as feelings [14,15] (Fig 1). After an emotional response, the discrepancies between anticipated outcomes and actual situations are integrated into future predictive models that correct novel mental representations and emotional states. This concept is pivotal in understanding how emotions are formed and regulated over time [9,12,14]. Teachers’ and students’ emotions may be related to past academic life events and future expectations. Since emotions can be regulated over time, they present ideal targets for educational interventions. Consequently, educators must understand the origins of emotions and how to manage them predictably.

**Fig 1.**
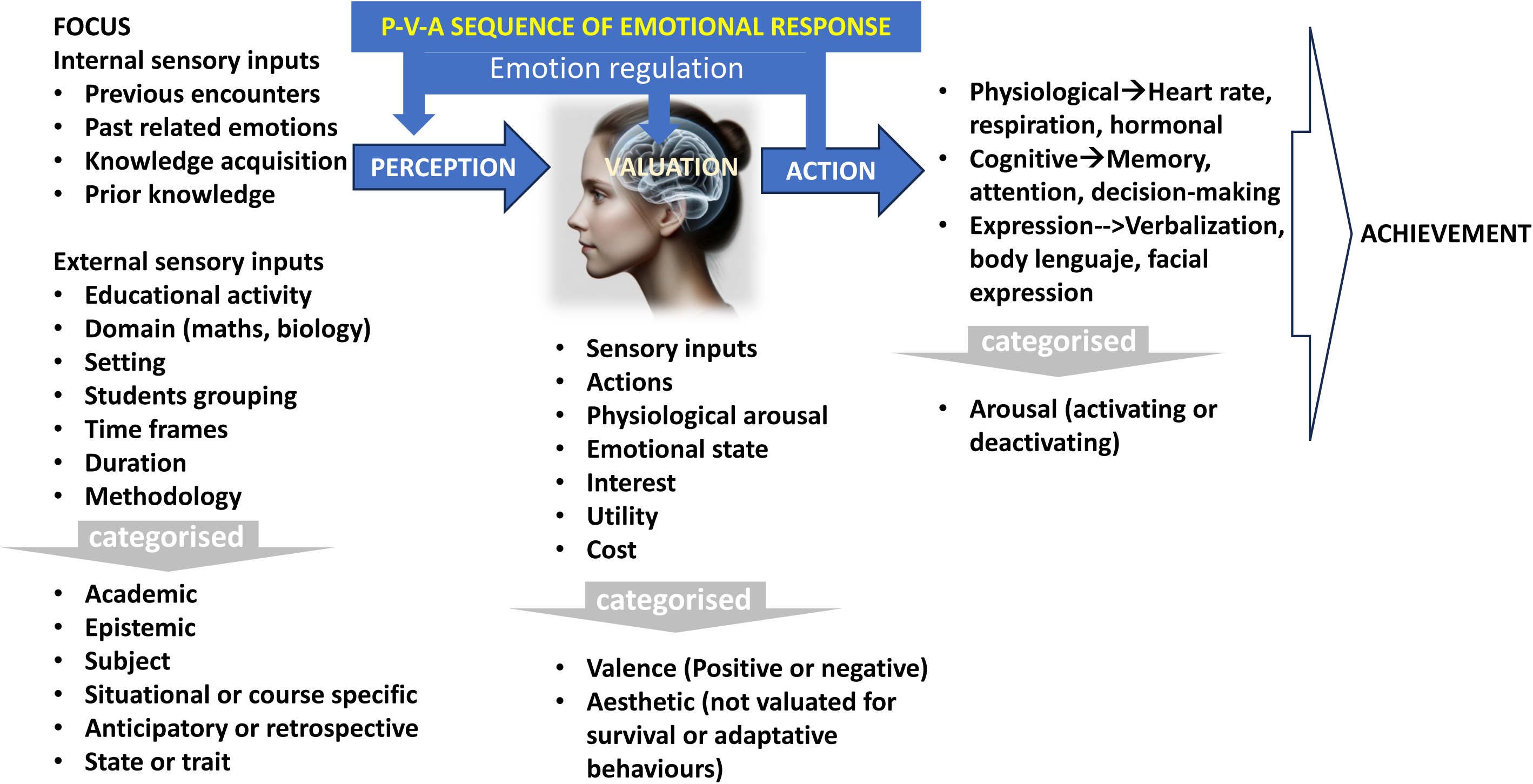
Categorisation of the elements involved in emotional reactivity (Perception-Valuation-Action sequence) and emotion regulation.

### 1.3 Categorising and classifying emotions

Based on the P-V-A sequence, emotions are categorised according to the perceived focus (object or source of stimulus), the valuation conducted, and the resulting action [16]. While not exhaustive, if the focus is an educational activity or its achievements, these are academic emotions (such as enjoyment or pride). If they stem from knowledge acquisition, they are epistemic emotions (such as surprise, curiosity, or confusion). These may further pertain to distinct domains (e.g., Language, Mathematics, or Science), contexts (e.g., laboratory, outdoor, or classrooms), time frames (situational or course-specific), time relations (anticipatory or retrospective), duration of the emotion (state or trait), methodologies (e.g., active learning or teacher-centred), and so forth. Valuation considers sensory inputs, actions, physiological arousal, emotional state [15], competence, unexpectedness, and subjective value [17]. According to these appraisals, emotions are qualified as utilitarian (if their utility value is considered) or aesthetic emotions (derived from music or art, where utility is not a consideration). Emotions are also divided according to their valence: positive if the value prediction is that it will be beneficial for the individual, and negative if the prediction is adverse. Regarding action, emotions are classified as activating (high arousal) if they activate the physiological system (such as enjoyment) or deactivating (low arousal) if they inhibit action (such as boredom). The absence of a consistent and comprehensive categorisation of emotions in numerous studies compromises their comparability. Expressing a singular emotion, such as enjoyment, receives varied interpretations contingent upon the P-V-A sequence. For example, in academic settings, enjoyment may be induced by various stimuli, including acquiring knowledge, overcoming challenges, or a preference for specific subjects. While enjoyment is generally perceived as a manifestation of attraction towards stimulus (positive valence) and is associated with motivation in educational environments, it can also be counterproductive (negative value), as a distraction, diminishing academic achievement [18]. Nervousness towards mathematics as a whole is considered inhibiting, yet towards a specific problem, it could be activating [18]. Joy can be classified as either outcome prospective or retrospective. In other words, to achieve generalizability in emotion research and to facilitate the development of viable emotion regulation strategies, it is crucial to contextualise the focus of emotions, the appraisals involved, and the potential resultant actions. This contextualisation is more precise in immediate or situational events than past events or course-specific studies, commonly used to explore the interplay of emotions and learning [17].

### 1.4 Emotions and the learning process

Both psychological and neurophysiological studies concur that emotions are intrinsically linked to students’ learning processes and outcomes. Emotions govern attention, shape motivation to learn, and alter learning strategies, thus enabling a more focused engagement with pertinent information. Emotions influence human cognitive processes, encompassing perception, learning, memory, reasoning, episodic recall, cognitive control processes, decision-making, spontaneous thought, and problem-solving [10,18,19]. Furthermore, emotions facilitate long-term learning since emotionally charged information is more readily recalled than neutral information [10,18]. These insights support the idea that emotion operates as a fundamental feature of cognition. Emotion regulates the flow of information across brain regions and, in conjunction with a suite of neuromodulators, ensures that emotional outcomes are the focal point of perception, thought, and action [10,19]. Emotion dynamics such as valence shifts transform otherwise neutral experiences into meaningful, memorable events [20]. Given that the dynamics of emotional states shape the episodic structure of memory [20], it is essential to examine the interplay between emotions and learning within the classroom context, in situ, where shifts in valence and arousal occur.

There is broad consensus on the significant role emotions play in academic achievement. Emotions such as enjoyment, boredom, and anxiety can profoundly impact students’ learning experiences and outcomes [5,17,21,22]. This influence is observed both in situational contexts (course-specific activities) and across broader domain-wide contexts, though the effects may vary depending on the setting. Classroom activities, assignments, or immediate educational challenges evoke situational emotions directly affecting performance [23]. Among preservice teachers, science activities such as geology, chemistry, microbiology, and cell biology involve significant emotion regulation, enhancing positive emotions (e.g., joy, enjoyment, enthusiasm) during the practice and reducing negative emotions (e.g., worry, frustration, nervousness). In these activities, both anticipatory emotions and emotions experienced during the practices are closely associated with academic achievement [24–28]. Emotions like frustration, boredom, or enjoyment are closely tied to specific classroom tasks, and their impact on performance is more immediate and short-term [23]. In contrast, domain-wide emotions, such as math anxiety, are linked to general subject areas like mathematics or science and may persist over time, influencing performance across multiple courses within the same domain and differing significantly between subjects [21,29].

Boredom has a significant effect on learning. As a negative deactivating emotion, it can reduce attention and motivation, often resulting in disengagement from the learning process. The reciprocal relationship between boredom and academic achievement has been repeatedly documented, showing its negative impact across various contexts and subject domains [5,30–32]. Boredom is considered an adaptive response that regulates an individual’s effort in a specific situation [33]. Within the framework of Control-Value Theory, boredom is associated with low levels of perceived control and low value placed on the task at hand, further contributing to reduced engagement and performance [34]. Achievement-related subjective control and value negatively predict boredom [35]. Additionally, beliefs about prospective control—rather than current perceived control—may better predict engagement and boredom [36]. The mere expectation that a lecture will be boring can intensify the actual experience of boredom [37]. This suggests that discrepancies between anticipated and actual control may exacerbate boredom, with emotion regulation playing a role in adjusting emotional states based on these discrepancies. These observations may have important educational implications, as addressing boredom in the first session of a subject has been shown to improve achievement in subsequent sessions [38].

In addition to context, subject, and domain, the variability of discrete emotions such as anxiety, enjoyment, and boredom can also be explained by sex and prior knowledge. Females tend to experience higher levels of anxiety in academic settings, particularly during assessments. However, these differences are less pronounced for other emotions, such as enjoyment and boredom [31,39,40]. Students with more domain-specific knowledge tend to experience more positive emotions, such as enjoyment, and fewer negative emotions, such as frustration and boredom, particularly during challenging activities [41,42].

However, Most research on the interplay between emotions and learning has focused on overall course performance rather than specific learning situations or methodologies [17]. It is necessary to analyse the role of emotions in learning specific scientific content by assessing emotions and learning using temporally and situationally circumscribed measures. Such an approach would facilitate comparing and generalising research findings across various situational contexts. However, there is a substantial lack of studies conducted in real-life classroom settings [17,43,44]. Two main factors contribute to this research gap: firstly, the wide variety of domains, methodologies, and classroom settings; secondly, the absence of validated instruments to accurately assess emotions in course-specific, situational contexts.

### 1.5 Assessing emotions

Although emotions are intrinsically personal (encompassing past experiences) and are variable processes that are situational, they manifest commonalities when individuals communicate their feelings, revealing variances across cultural contexts [17]. Emotions are often measured using self-report questionnaires [45]. While these only capture a subset of the action programs elicited by human emotions [11], they are associated with specific brain region activation, thereby bridging neurophysiological and psychological findings [10,46]. Emotions can be analysed both as discrete categories (e.g., joy and boredom) and as dimensions our brains use different representational dynamics that emphasise either categories or dimensions [47]. Unlike discrete emotions, latent factors tend to exhibit less variability and may depend on different moderators. A recent study explored within-person and between-person variance in discrete emotions and latent integrative affect factors in relation to school performance. The findings suggest that discrete emotions display higher contextual variability, while latent factors are more stable and influenced by distinct moderators [48].

In the context of Control-Value Theory, the Achievement Emotions Questionnaire (AEQ) stands out as a validated instrument for assessing eight emotions (6 to 12 items per scale) linked to academic activities and achievements, with its briefest version (4 items per scale) taking 15 minutes to administer [49]. While effective for tracking trait emotions over entire courses, it is less operative at capturing situational emotions within specific activities of less than one hour, contributing to the scarcity of situational emotions research [50]. Furthermore, some emotions are often overlooked in this instrument. For instance, disgust is scarcely addressed within the framework of Control-Value Theory, as, according to Pekrun [13] "There was virtually no major human emotion not reported by our participants, with disgust being the only notable exception”.

Assessing situational (state) emotions requires validated experience sampling methods and instruments that can be rapidly administered without detriment to the ongoing activity or the assessment of situational emotions [44]. Recent developments have led to less intrusive methods, including facial recognition [51] and abbreviated questionnaires that preserve more extended versions’ reliability and significant correlations [50,52,53]. Accumulated evidence supports the equivalence in reliability and validity between single-item and multi-item measures at the school level, providing teachers with information to orient their teaching practice [52,53]. Although single-item Likert scale ratings are commonly employed in emotion measurement, existing instruments often lack measurement invariance across different cohorts [54,55]. While single-item instruments are practical and widely used to capture situational, course-specific emotions, there remains a significant gap in the availability of validated instruments tailored to the diverse needs of practitioners across different educational contexts, which hampers the broader application of emotional assessments in everyday classroom settings.

We have recently designed and used a short instrument for assessing ten emotions via single-item measures. This instrument captures the emotional differences of students toward science lectures and practical sessions [56]. Furthermore, it has been utilised to examine shifts in emotions before and after an educational intervention in microbiology designed for preservice teachers [57]. This instrument enables the extraction of two latent factors to explore the shared variance among five activating positive and four activating negative emotions, the analysis of their correlations with learning outcomes and the construction of a robust network of emotions and their influence on academic performance [58]. The brief completion time of less than five minutes markedly minimised classroom disruptions. The instrument showed internal and external validity when measuring anticipatory emotions. Our previous findings corroborated earlier research on the link between emotions and academic achievement and showed that students lacking sufficient prior knowledge and those pursuing non-scientific upper-secondary education programs tend to anticipate more negative emotions. These emotions, in turn, were linked to subsequent learning, albeit with varying impacts: while frustration (a dampening emotion) hindered learning, nervousness (a stimulating emotional state) appeared to facilitate it [57,58].

### 1.6 This study

This study aims to carry out a conceptual replication of our previous study [57], substituting the microbiology lab experience with DNA extraction using household materials. In addition, we have expanded the discrete emotions measured by the instrument from ten to sixteen and have further validated and established the measurement invariance of the two latent factors: positive and negative activating emotions. Our findings reaffirm the regulatory function of emotions on achievement outcomes, with boredom emerging as a particularly influential factor. The observed differences between the two interventions—microbiology lab experience and DNA extraction—suggest that emotion regulations in classroom activities should be addressed on a case-by-case basis. This work contributes to a deeper understanding of the role of anticipatory situational emotions in education.

## 2. Methodology

### 2.1 Sample

This study involved a non-probabilistic sample of 269 preservice teachers (average age of 21 years, 62% female) pursuing a primary education degree at the University of Extremadura on two campuses (Cáceres and Badajoz) of the same Spanish region. All participants were enrolled in the Earth and Life Science Education course during Semester 6, across two consecutive academic years, a course that introduced them to the fundamentals of biology education. Concerning their prior education, 72% had a non-scientific Upper-Secondary Education degree (coded as non-STEM) in humanities, social sciences, or arts, while the remaining 28% held a scientific one (coded as STEM) (Table 1). The participant count exceeds the minimum required number of 262, as estimated using G*power.

**Table 1.** Participants distributed by sex and prior education during the upper secondary education itinerary (STEM or non-STEM).

| Sex | Prior education | N |
| --- | --- | --- |
| Females | STEM | 37 |
|  | Non-STEM | 130 |
| Males | STEM | 37 |
|  | Non-STEM | 65 |
| TOTAL |  | **269** |

### 2.2 Procedure

Participants engaged in an experimental activity related to DNA extraction, implemented through guided inquiry. Briefly, this activity shows fundamental cell biology, physics, and chemistry concepts while highlighting science-technology-society interactions through various biotechnology applications. The teacher facilitated students’ understanding of DNA extraction principles using everyday materials, guiding them through various problems, questions, and debates. Participants then developed and applied their extraction protocols to plant (tomato) and animal (saliva) samples (Fig 2). This activity has been validated and shown to be effective for active interdisciplinary science learning [28].

**Fig 2.**
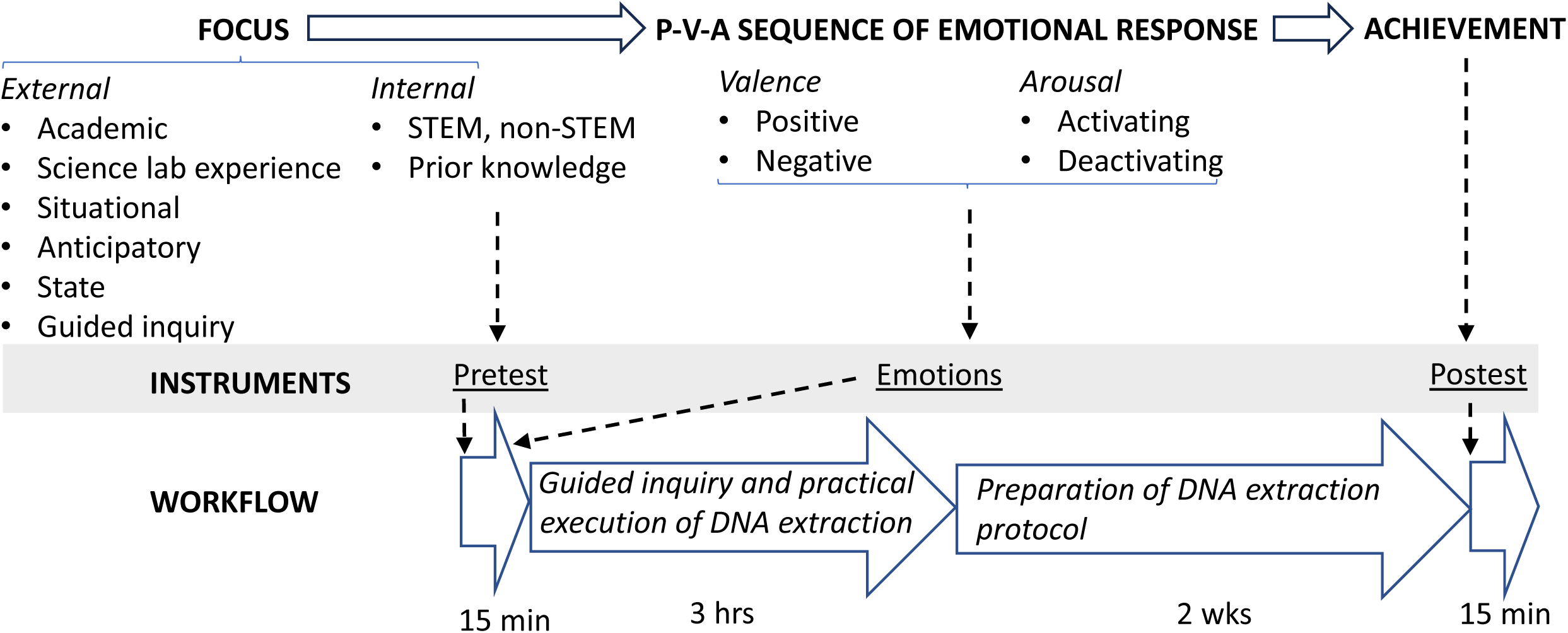
Elements of emotional reactivity relevant to this study, instruments, and workflow integration. The top panel presents the attributes used to categorise emotions based on focus, emotional response, and their relationship to achievement. Detailed descriptions of these emotions are provided in Table 1. Dotted lines indicate the connections between prior knowledge, emotions, and achievement, along with the instruments used to measure each (grey background). These connections extend to the application of the instruments within the workflow timeline, shown in the bottom panel.

Before the activity commenced, participants were fully informed. They provided their written informed consent per the statement below: ‘You have been invited to participate in a research study to improve teaching at the University. The study seeks to understand the relationship between learning and the student’s perception of the activities carried out in the classroom. In this case, DNA extraction with everyday materials. In future publications, the data obtained will be presented globally, fully preserving the individual characteristics of the participants, who will always remain anonymous. Participation is voluntary, poses no risk, and does not affect the participants’ grades in any way. The project lasts for three years and is not for profit. The participant may withdraw at any time.’ (Supplementary Material)."All participants completed a pretest questionnaire assessing anticipatory emotions. After this, 192 participants completed a second questionnaire on scientific knowledge. In contrast, a separate subset of 77 participants did not complete the second questionnaire, serving as a control group to examine the effect of pretest completion on learning outcomes. The emotions questionnaire was administered first to prevent knowledge questionnaire resolution from interfering with anticipatory emotions. Fifteen days post-intervention, students submitted a report on the practice as part of the intervention, and 255 participants completed a posttest assessing subsequent knowledge (Fig 2). The study was approved by the University of Extremadura’s Ethics Committee (registry 185/2021) during its session held on December 15, 2021.

### 2.3 Instruments

Anticipatory emotions were assessed through an extended version of a previously validated quantitative self-report questionnaire [56]. This questionnaire was selected for its quick administration and reliability. Its structure is similar to the frequently used PANAS instrument, which contains two 10-item mood scales, allowing the extraction of two scales [59]. Our instrument scrutinised sixteen academic emotions and associated states that could potentially influence the course of the educational activity. This encompassed six positive activating emotions (joy, enjoyment, enthusiasm, gratitude, pride and awe), six negative activating emotions (worry, frustration, nervousness, shame, fear and disgust), two positive deactivating emotions (confidence, and satisfaction), one negative deactivating emotions (boredom) and a cognitive state characterised as uncertainty. All the aforementioned items are conceptualised within the Control-Value Theory of Achievement (Table 2). These emotions combine different dimensions of Pekrun’s taxonomy [17]: object focus (activity and outcome), valence (positive and negative), and arousal (activating and deactivating). The positive activating emotions (joy, enjoyment, enthusiasm, gratitude, pride, and awe) and the negative activating emotions (worry, frustration, nervousness, shame, fear, and disgust) were used to extract two latent factors (Fig 3), one for each valence: PAE (Positive activating emotions) and NAE (Negative activating emotions). Participants were prompted with the following inquiry: "What emotions do you anticipate experiencing when confronted with a DNA extraction practice using everyday materials?" This question effectively captured the subjective aspect of emotional experience, the temporal dimension (future emotions), and the specific object focus (the activity of DNA extraction with everyday material). All items were answered using a Likert-scale ranging from (1) not at all to (5) very much. At the onset of the questionnaire, participants provided information regarding their sex and prior educational background. Additionally, they generated an anonymous code to pair their pretest and posttest responses.

**Fig 3.**
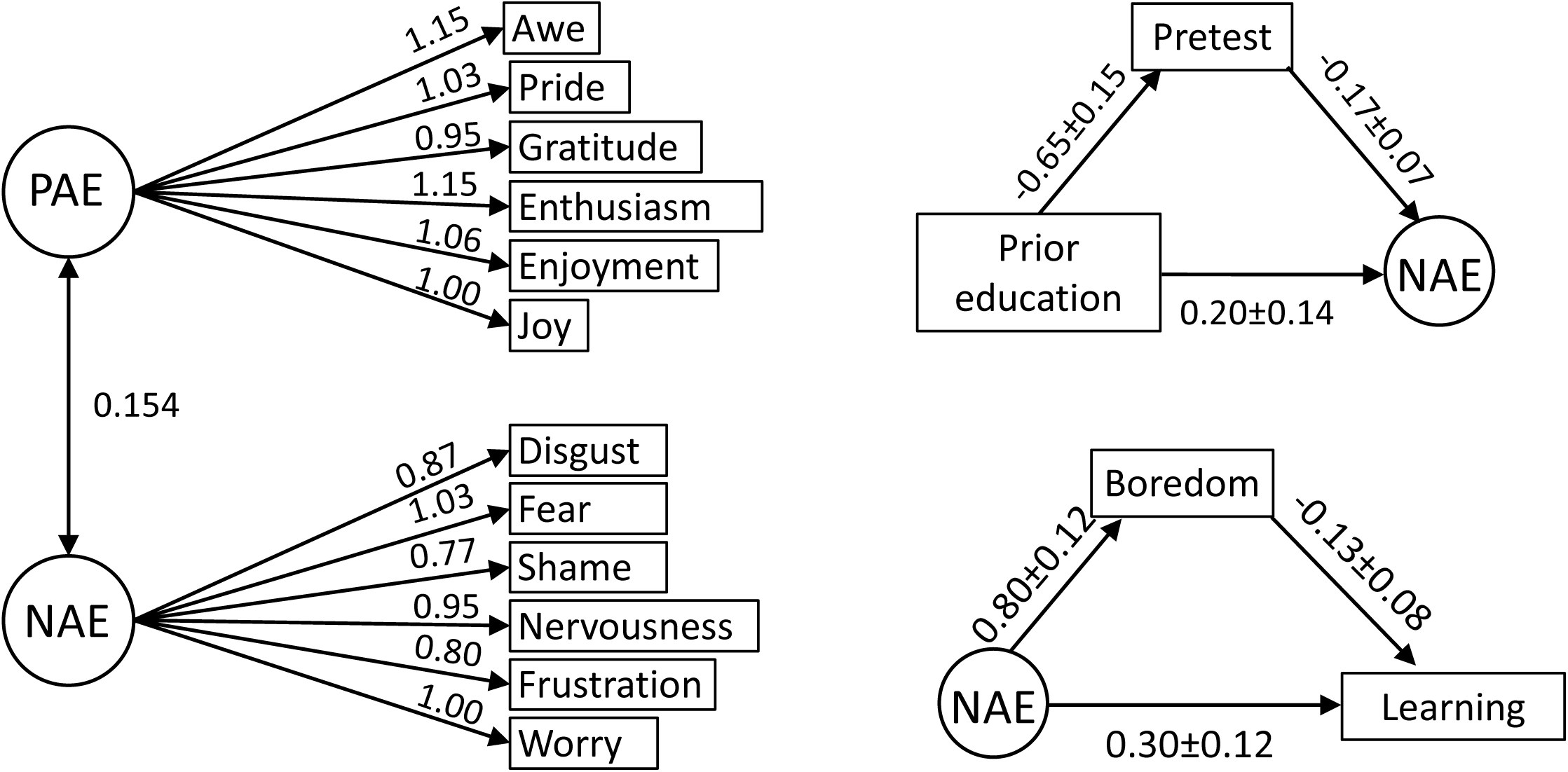
Analytical models for factors of activating emotions and mediation effects. Left: Factor analysis model for positive (PAE) and negative (NAE) activating emotions scales. The numbers on the uni-directional arrows represent standardised coefficients, which indicate the strength and influence of each factor on the associated emotions. The numbers on the double-headed arrows denote Spearman factor correlations. Right: The diagrams show mediation models for pretest (top) and boredom (bottom). Numbers in the mediation models represent path coefficients and their standard errors. Observable variables are represented in boxes, whereas latent variables are represented in circles.

**Table 2.**
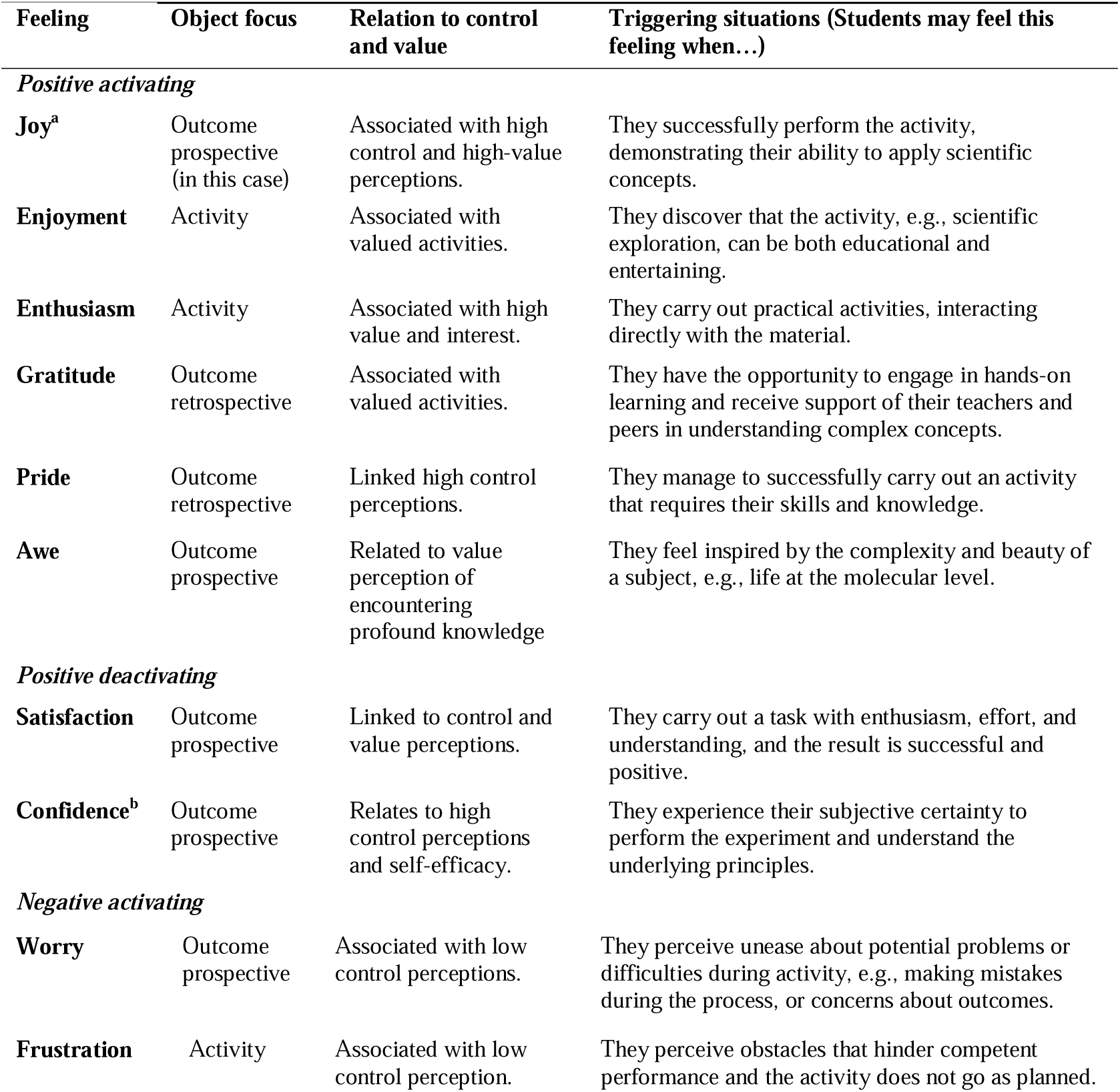
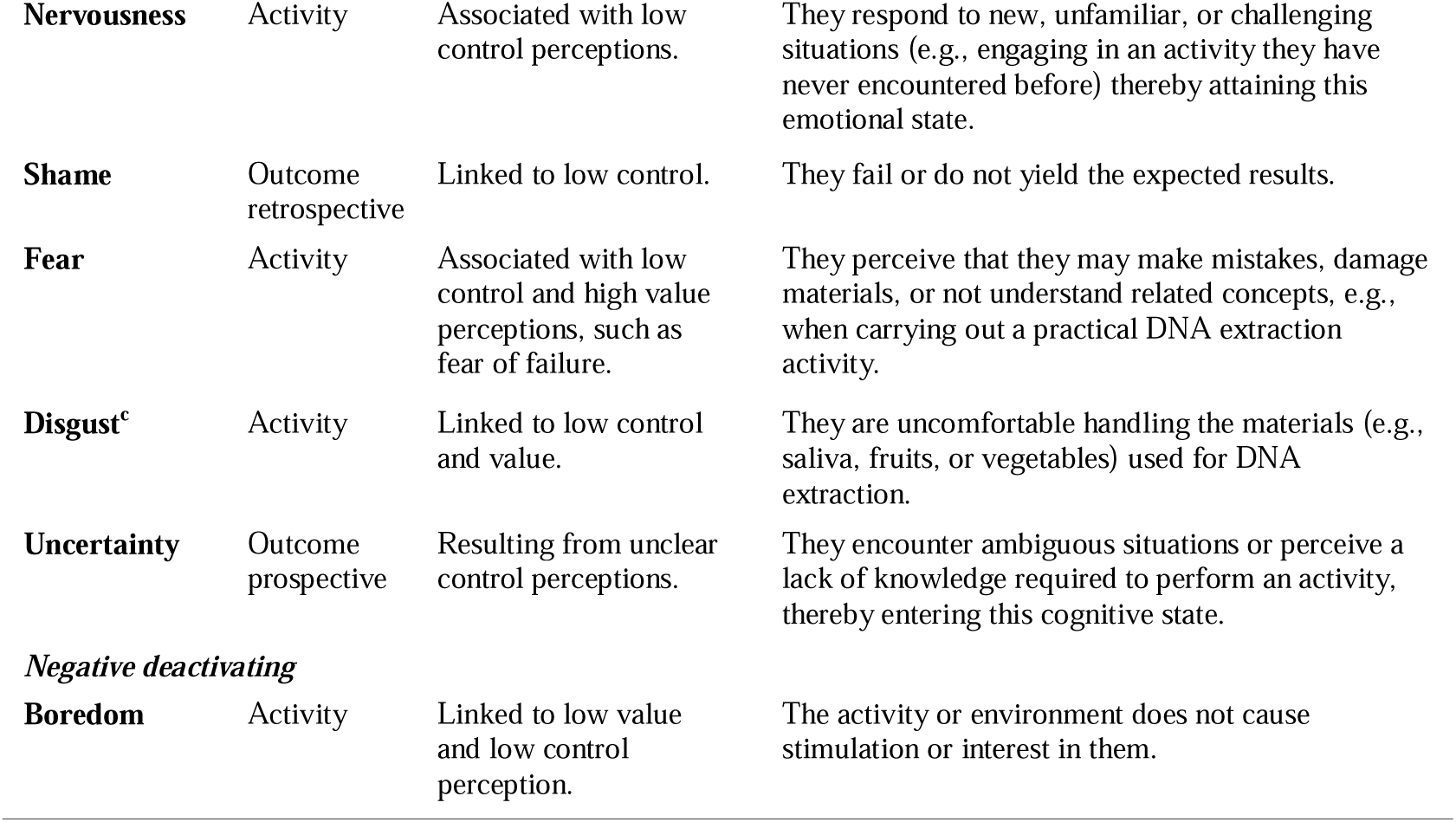
Characteristics and supposed triggering situation of feelings (as expression of the emotions, emotional responses, and cognitive states) involved in the study.

| Feeling | Object focus | Relation to control and value | Triggering situations (Students may feel this feeling when...) |
| --- | --- | --- | --- |
| *Positive activating* |  |  |  |
| **Joy<sup>a</sup>** | Outcome prospective (in this case) | Associated with high control and high-value perceptions. | They successfully perform the activity, demonstrating their ability to apply scientific concepts. |
| **Enjoyment** | Activity | Associated with valued activities. | They discover that the activity, e.g., scientific exploration, can be both educational and entertaining. |
| **Enthusiasm** | Activity | Associated with high value and interest. | They carry out practical activities, interacting directly with the material. |
| **Gratitude** | Outcome retrospective | Associated with valued activities. | They have the opportunity to engage in hands-on learning and receive support of their teachers and peers in understanding complex concepts. |
| **Pride** | Outcome retrospective | Linked high control perceptions. | They manage to successfully carry out an activity that requires their skills and knowledge. |
| **Awe** | Outcome prospective | Related to value perception of encountering profound knowledge | They feel inspired by the complexity and beauty of a subject, e.g., life at the molecular level. |
| *Positive deactivating* |  |  |  |
| **Satisfaction** | Outcome prospective | Linked to control and value perceptions. | They carry out a task with enthusiasm, effort, and understanding, and the result is successful and positive. |
| **Confidence<sup>b</sup>** | Outcome prospective | Relates to high control perceptions and self-efficacy. | They experience their subjective certainty to perform the experiment and understand the underlying principles. |
| *Negative activating* |  |  |  |
| **Worry** | Outcome prospective | Associated with low control perceptions. | They perceive unease about potential problems or difficulties during activity, e.g., making mistakes during the process, or concerns about outcomes. |
| **Frustration** | Activity | Associated with low control perception. | They perceive obstacles that hinder competent performance and the activity does not go as planned. |
| **Nervousness** | Activity | Associated with low control perceptions. | They respond to new, unfamiliar, or challenging situations (e.g., engaging in an activity they have never encountered before) thereby attaining this emotional state. |
| **Shame** | Outcome retrospective | Linked to low control. | They fail or do not yield the expected results. |
| **Fear** | Activity | Associated with low control and high value perceptions, such as fear of failure. | They perceive that they may make mistakes, damage materials, or not understand related concepts, e.g., when carrying out a practical DNA extraction activity. |
| **Disgust<sup>c</sup>** | Activity | Linked to low control and value. | They are uncomfortable handling the materials (e.g., saliva, fruits, or vegetables) used for DNA extraction. |
| **Uncertainty** | Outcome prospective | Resulting from unclear control perceptions. | They encounter ambiguous situations or perceive a lack of knowledge required to perform an activity, thereby entering this cognitive state. |
| *Negative deactivating* |  |  |  |
| **Boredom** | Activity | Linked to low value and low control perception. | The activity or environment does not cause stimulation or interest in them. |

Scientific knowledge was assessed using a validated questionnaire with closed-ended test-type questions. The questions addressed common misconceptions among secondary school students and preservice teachers [60] and were based on key concepts from the Trends in International Mathematics and Science Study report [61]. Learning was assessed as a posttest score minus a pretest score.

### 2.4 Data analysis

We employed three distinct approaches for data analysis: the analysis of single items of feelings, factor analysis, and network analysis. Examining individual items preserves item-specific information, though it can lack structure and become challenging to interpret when many variables are involved. Factor analysis is effective for grouping variables and identifying latent traits, facilitating the interpretation of larger datasets, though it may overlook the finer details of individual items. Network analysis provides valuable insights into the interactions between items, offering a systemic view of their relationships. However, it can be complex and, unlike factor analysis, does not identify unobserved variables that drive these relationships. All descriptive and statistical analyses were conducted using Jamovi 2.48 [62] and JASP 0.18.1 [63,64]. We used G*power 3.1.9.6 [65] for sample size calculation and statistical power analysis (α = 0.05, Power = 0.80, Effect size = 0.3).

According to the Shapiro-Wilk test, the data did not follow a normal distribution (Table 3). Non-parametric statistical methods were employed, including Spearman correlation for variable interactions and Robust ANOVA to discern sources of variability in unbalanced subgroups lacking normality. For Robust ANOVA, we employed the Walrus module of Jamovi [66], using a trimmed means level of 0.2.

**Table 3.**
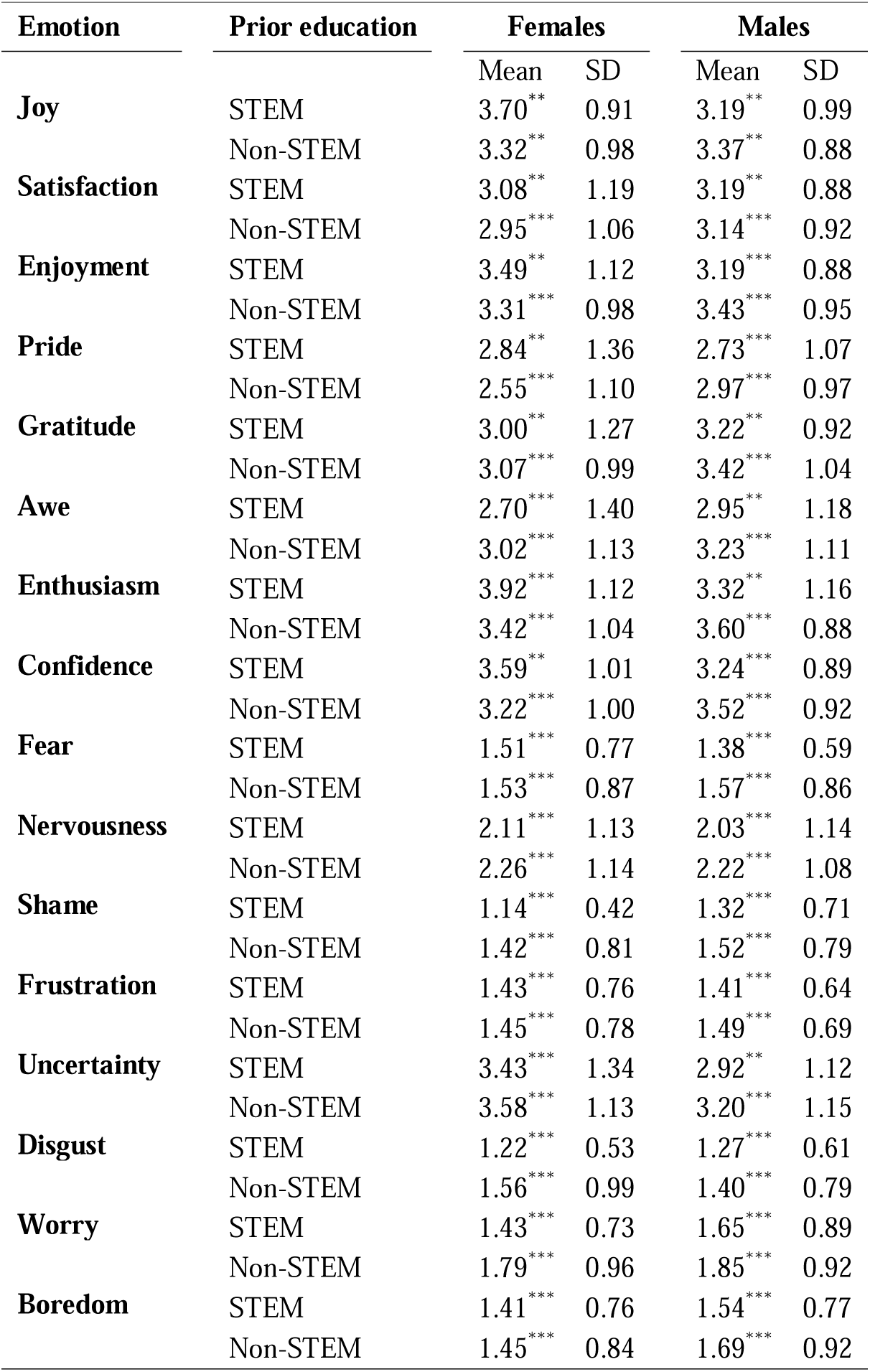
Distribution of feelings reported by participants according to sex and prior education during the upper secondary education itinerary (STEM or non-STEM). P-values of the Shapiro-Wilk normality test, **<0.01, ***<0.001.

| Emotion | Prior education | Females |  | Males |  |
| --- | --- | --- | --- | --- | --- |
|  |  | Mean | SD | Mean | SD |
| **Joy** | STEM | 3.70** | 0.91 | 3.19** | 0.99 |
|  | Non-STEM | 3.32** | 0.98 | 3.37** | 0.88 |
| **Satisfaction** | STEM | 3.08** | 1.19 | 3.19** | 0.88 |
|  | Non-STEM | 2.95*** | 1.06 | 3.14*** | 0.92 |
| **Enjoyment** | STEM | 3.49** | 1.12 | 3.19*** | 0.88 |
|  | Non-STEM | 3.31*** | 0.98 | 3.43*** | 0.95 |
| **Pride** | STEM | 2.84** | 1.36 | 2.73*** | 1.07 |
|  | Non-STEM | 2.55*** | 1.10 | 2.97*** | 0.97 |
| **Gratitude** | STEM | 3.00** | 1.27 | 3.22** | 0.92 |
|  | Non-STEM | 3.07*** | 0.99 | 3.42*** | 1.04 |
| **Awe** | STEM | 2.70*** | 1.40 | 2.95** | 1.18 |
|  | Non-STEM | 3.02*** | 1.13 | 3.23*** | 1.11 |
| **Enthusiasm** | STEM | 3.92*** | 1.12 | 3.32** | 1.16 |
|  | Non-STEM | 3.42*** | 1.04 | 3.60*** | 0.88 |
| **Confidence** | STEM | 3.59** | 1.01 | 3.24*** | 0.89 |
|  | Non-STEM | 3.22*** | 1.00 | 3.52*** | 0.92 |
| **Fear** | STEM | 1.51*** | 0.77 | 1.38*** | 0.59 |
|  | Non-STEM | 1.53*** | 0.87 | 1.57*** | 0.86 |
| **Nervousness** | STEM | 2.11*** | 1.13 | 2.03*** | 1.14 |
|  | Non-STEM | 2.26*** | 1.14 | 2.22*** | 1.08 |
| **Shame** | STEM | 1.14*** | 0.42 | 1.32*** | 0.71 |
|  | Non-STEM | 1.42*** | 0.81 | 1.52*** | 0.79 |
| **Frustration** | STEM | 1.43*** | 0.76 | 1.41*** | 0.64 |
|  | Non-STEM | 1.45*** | 0.78 | 1.49*** | 0.69 |
| **Uncertainty** | STEM | 3.43*** | 1.34 | 2.92** | 1.12 |
|  | Non-STEM | 3.58*** | 1.13 | 3.20*** | 1.15 |
| **Disgust** | STEM | 1.22*** | 0.53 | 1.27*** | 0.61 |
|  | Non-STEM | 1.56*** | 0.99 | 1.40*** | 0.79 |
| **Worry** | STEM | 1.43*** | 0.73 | 1.65*** | 0.89 |
|  | Non-STEM | 1.79*** | 0.96 | 1.85*** | 0.92 |
| **Boredom** | STEM | 1.41*** | 0.76 | 1.54*** | 0.77 |
|  | Non-STEM | 1.45*** | 0.84 | 1.69*** | 0.92 |

We used a confirmatory factor analysis (CFA), as implemented in the structural equation modelling module of JASP, to confirm the presence of two underlying latent factors (PAE and NAE) behind the subset of six positive and six negative activating emotions (Fig 3A). While CFA is the most suitable and widely used method for factor extraction, it assumes normally distributed data, a condition not met in this case. We used an Oblimin rotation [56], which is suitable for correlated factors. Since there is no agreement about the best model fit index for CFA of categorical data [67], we provide various fit indices and thresholds recommended to assess the adequacy of the model when using categorical data [67–69]. These model fit indexes include χ² Test, Root Mean Square Error of Approximation (RMSEA), Tucker-Lewis Index (TLI), and Comparative Fit Index (CFI). χ² Test is sensitive to sample size, and it is shown for comparison. Acceptable fit thresholds are >0.05 for the χ² test, <0.08 for RMSEA, and >0.90 for both TLI and CFI [67–69]. Other factor metrics, including MacDonald’s ω (composite reliability for internal consistency), AVE (Average Variance Extracted), and HTMT (Heterotrait-Monotrait ratio of correlations), were calculated using JASP software. PAE and NAE factor scores were saved from CFA.

We use measurement invariance (MI) analysis to compare the equivalence of measures across groups (sex and prior education). We used CFA to define a sequence of nested models. Configural invariance was tested by fitting the two-factor model structure to different groups to see if the same general factor structure holds. Metric invariance was tested by constraining factor loadings to be equal across groups. Scalar invariance was tested by additionally constraining item intercepts to be equal across groups. The changes in χ² (Δχ²), RMSEA (ΔRMSEA), and CFI (ΔCFI) across the nested models were used to evaluate whether the construct was invariant at each of the stages of the multigroup analysis mentioned above. As recently recommended, a scale was considered non-invariant with respect to thresholds if both a significant χ^2^ and at least one between a RMSEA > .01 or a CFI < −.01 was found [69]. Statistical mediation and moderation analyses [70] were conducted using the Medmod module of Jamovi. Moderation analysis employs multiple regression to determine the impact of each variable and its interaction with other variables. For these analyses, the values of R^2^ (proportion of explained variance), β (regression coefficient or relationship), and p (probability of observed results occurring by chance) were determined. These analyses are valuable tools for understanding mechanisms behind observed relationships. However, they rely heavily on appropriate model specification, accurate measurement, and the assumption of no confounding variables (Fig 3B).

We used a network analysis (Network module of JASP) to estimate the graphical structure of a network of variables, within the framework of covariance selection models (EBICglasso estimator). The EBICglasso estimator produces simple models and mitigates overfitting, especially in high-dimensional data. Its key merit lies in providing robust estimates even with small sample sizes. However, it may lead to underfitting if the true model is more complex. The resulting network represents variables as nodes and their interactions as edges. Two variables were connected if they correlated and shared the maximum unique covariance [71].

## 3. Results and discussion

We analyse the predictive capacity of emotions in fostering learning and academic achievement. To eliminate the possibility that variations in achievement stemmed from either teacher effectiveness or the learning gained during the completion of the pretest [72], we initially confirmed that the posttest scores were independent of both the participants’ campus (each with a different instructor, Mann-Whitney U test p-value 0.52) and the completion of the pretest (Mann-Whitney U test p-value 0.24). As is common in the demographic composition observed in educational degree programs, the cohort under study exhibits a demographic skew, with a preponderance of females and participants with a non-scientific orientation (non-STEM) during their upper secondary education tenure (Table 1).

### 3.1 Variability of discrete emotions

Participants were administered a questionnaire wherein they self-reported their feelings on a Likert scale. Variability in participants’ anticipatory emotions was observed, showing deviations from normal distribution for sex and prior education (Table 3). Considering this and the unbalance in sample size of subgroups (Table 1), we employed Robust ANOVA analysis to discern the sources of variability (Table 4). Concerning sex, our robust ANOVA analysis revealed that male participants exhibited greater expressions of gratitude (p < 0.05) compared to their female counterparts, alongside lower levels of uncertainty (p < 0.01). This contrasts with findings from a microbiology activity [58], where males exhibited greater confidence than females, pointing out that gender differences are likely specific to each science activity. These results align with previous research indicating gender differences in the expression of emotions, e.g., female preservice teachers often have more negative and less positive emotions towards science than their male counterparts [22,73,74].

**Table 4.**
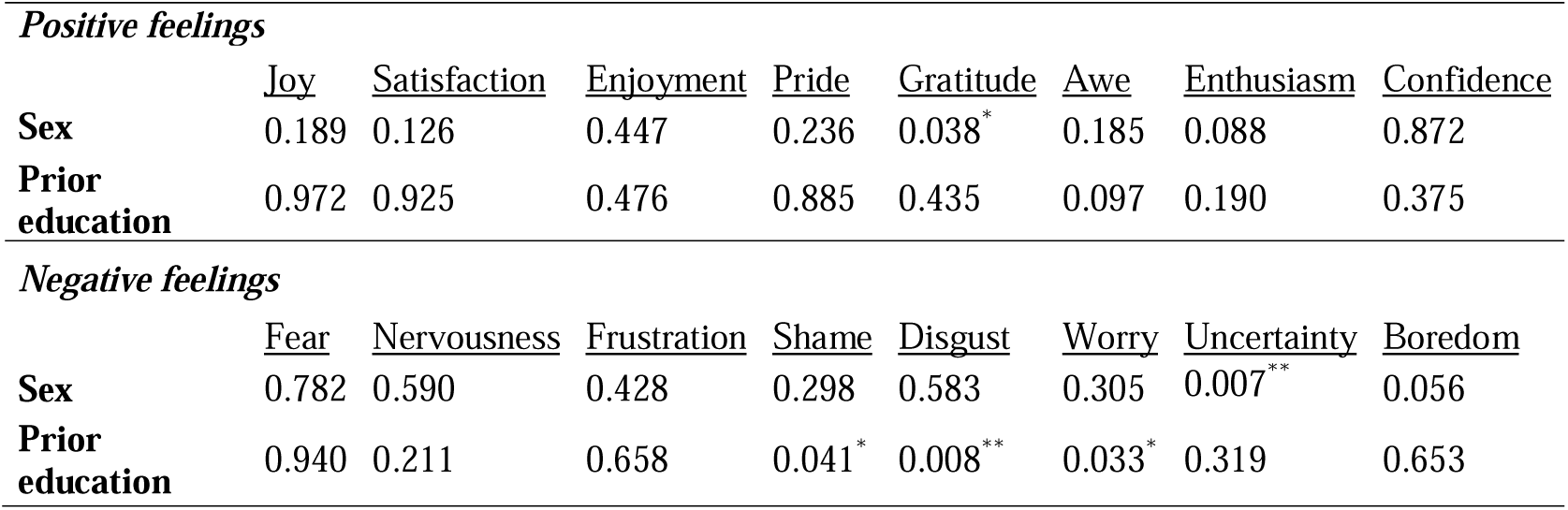
P-values (* <0.05, **<0.01) from robust ANOVA assessing the significance of sex and prior education in explaining the variability of positive (top) and negative (bottom) feelings.

| *Positive feelings* |  |  |  |  |  |  |  |  |
| --- | --- | --- | --- | --- | --- | --- | --- | --- |
|  | <u>Joy</u> | <u>Satisfaction</u> | <u>Enjoyment</u> | <u>Pride</u> | <u>Gratitude</u> | <u>Awe</u> | <u>Enthusiasm</u> | <u>Confidence</u> |
| **Sex** | 0.189 | 0.126 | 0.447 | 0.236 | 0.038* | 0.185 | 0.088 | 0.872 |
| **Prior education** | 0.972 | 0.925 | 0.476 | 0.885 | 0.435 | 0.097 | 0.190 | 0.375 |
| *Negative feelings* |  |  |  |  |  |  |  |  |
|  | <u>Fear</u> | <u>Nervousness</u> | <u>Frustration</u> | <u>Shame</u> | <u>Disgust</u> | <u>Worry</u> | <u>Uncertainty</u> | <u>Boredom</u> |
| **Sex** | 0.782 | 0.590 | 0.428 | 0.298 | 0.583 | 0.305 | 0.007** | 0.056 |
| **Prior education** | 0.940 | 0.211 | 0.658 | 0.041* | 0.008** | 0.033* | 0.319 | 0.653 |

Concerning the impact of prior education, participants with a scientific background during secondary education significantly anticipated fewer negative emotions, including shame, disgust, and worry, compared to their counterparts with a background in humanities, social sciences, or arts (Robust ANOVA, p < 0.05). This contrasts with findings from a microbiology activity [58], where a STEM prior education enhanced participants’ positive emotions (confidence and satisfaction) but not negative emotions. These disparities in anticipatory emotions may be attributed to variations in prior experiences, as evidenced by the higher scientific knowledge level (pretest score) among participants with a STEM background (Table 5). Such knowledge is known to have a lasting impact on emotional responses [40,56]. Our results confirm the impact of gender and prior education on anticipatory emotions [42] and point out that this impact is different for each science activity.

**Table 5.** P-values (***<0.001) from robust ANOVA assessing the significance of sex and prior education in explaining the variability of prior knowledge (pretest scores), achievement (posttest scores), and learning (posttest minus pretest).

|  | <u>Prior knowledge</u> | <u>Achievement</u> | <u>Learning</u> |
| --- | --- | --- | --- |
| **Sex** | 0.926 | 0.103 | 0.483 |
| **Prior education** | 0.001*** | 0.001*** | 0.757 |

Interaction (Sex\*Prior education) p-values are not shown due to non-significance (p > 0.05).

### 3.2 Variability of PAE and NAE scales

Confirmatory Factor Analysis (CFA) results corroborate the acceptability of the model, with model fit indices demonstrating satisfactory fit: RMSEA = 0.068 (< 0.08 cut-off), TLI = 0.920 (> 0.90 cut-off), and CFI = 0.935 (> 0.90 cut-off) (Table 6). High McDonald’s ω values (0.834 for PAE and 0.822 for NAE) support the internal consistency of the factors [75]. The McDonald’s ω composite reliability exceeding 0.6 validates the Average Variance Extracted (AVE) values for PAE and NAE (0.439 and 0.445, respectively) as acceptable [75], indicating a satisfactory variance capture relative to the variance attributed to measurement error. Moreover, the observed low Heterotrait-Monotrait (HTMT) ratio of correlations value of 0.142, substantially below the threshold of 0.85 [76], underscores robust discriminant validity, reinforcing the distinctiveness of the constructs (Table 6).

**Table 6.**
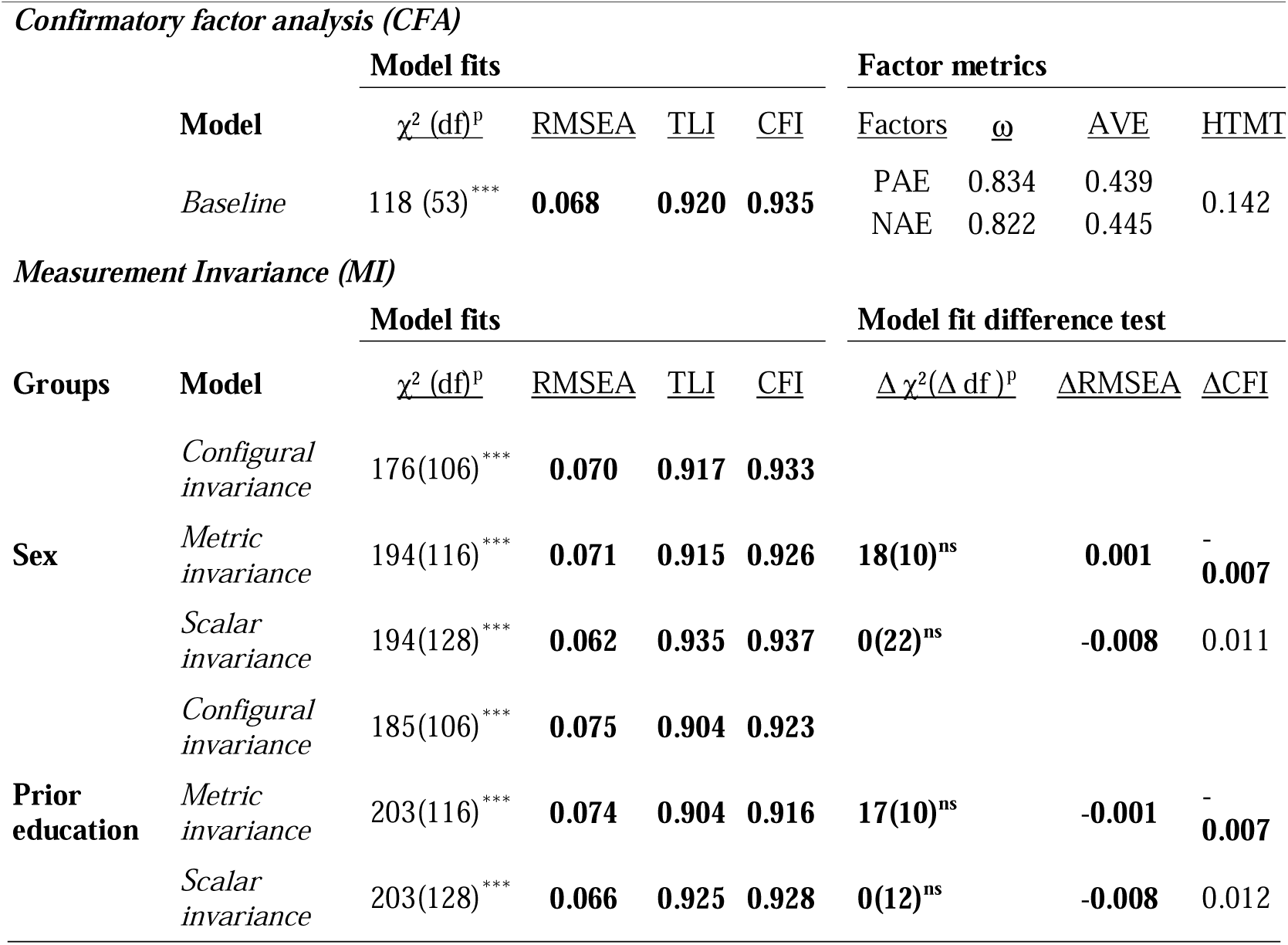
Model fit statistics for CFA of the two-factor model for positive activating (PAE) and negative activating (NAE) emotions scales (Top section), its MI across sex and prior education at each level of invariance testing, along with the changes in model fit (Bottom section) highlighting in bold the models that demonstrate acceptable fits (*** p-value<0.001; ns p-value>0.05).

Configural invariance analysis showed that this two-factor model was also acceptable for the sex (males and females) and prior education (STEM, and non-STEM) subgroups since the three model fit indexes are under (RMSE_sex_ 0.070<0.08; RMSE_prior-education_ 0.075<0.08) and over (TLI_sex_ 0.917>0.90; TLI_prior-education_ 0.935>0.90; CFI_sex_ 0.933>0.90; CFI_prior-education_ 0.923>0.90) the cut-off criteria. This two-factor model was also acceptable for all subgroups when loadings and intercepts were constrained in metric invariance and scalar invariance analysis, respectively. Moreover, model differences were under the cut-off criteria, demonstrating metric and scalar invariances. Upon progressing from configural to metric invariance analysis by introducing loading constraints, the change in model fit did not reach statistical significance as per the χ² difference test [Δ χ²__sex_(10) = 18, p = 0.056; Δ χ²__prior education_ (10) = 17, p = 0.070]. Concurrently, alternative fit measures supported acceptable changes: [ΔRMSEA__sex_ = 0.001 and ΔRMSEA__prior education_ = −0.001 (< 0.01 cutoff)] and [ΔCFI__sex_ = −0.007 and ΔCFI__prior education_ = −0.007 (< −0.01 cutoff)], thereby supporting the maintenance of metric invariance. Subsequent analysis, which incorporated intercept constraints, yielded an overall improvement in model fit without changes reaching statistical significance: [Δ χ²__sex_(22) = 0, p = 1; Δ χ²__prior education_ (12) = 0, p = 1], [ΔRMSEA__sex_ = −0.008 and ΔRMSEA__prior education_ = −0.008 (< 0.01 cutoff)] and [ΔCFI__sex_ = 0.011 and ΔCFI__prior education_ = −0.012 (< −0.01 cutoff)], thus confirming scalar invariance. Together, the CFA and measurement invariance analysis confirmed that the instrument provides a valid and reliable measure of latent constructs PAE and NAE, enabling unbiased comparisons across groups. In contrast to that observed with discrete emotions, the variability of factor scores is not explained by sex or prior education (Table 7). This finding aligns with previous research, which suggests that latent factors tend to be more stable and are influenced by different moderators [48]

**Table 7.** P-values from robust ANOVA assessing the significance of sex and prior education in explaining the variability of PAE and NAE factor scores.

|  | <u>PAE</u> | <u>NAE</u> |
| --- | --- | --- |
| Sex | 0.929 | 0.733 |
| Prior education | 0.846 | 0.170 |

27 Interaction (Sex\*Prior education) p-values are not shown due to non-significance ( $p > 0.05$ ).

Together, results support that the experience sampling method and instrument we used enabled us to collect near real-time data on individuals’ emotions. This method effectively minimises recall bias, which can occur when individuals access past emotional states, by capturing data at the moment it is experienced. In contrast, longer instruments like AEQ [22], typically used to assess emotions related to learning and performance over entire courses, are often retrospective and thus susceptible to recall bias. Students may struggle to accurately recall their emotions during specific activities, with responses potentially influenced by overall course outcomes rather than individual learning moments[44].

This methodology is particularly valuable for understanding dynamic, context-specific emotional phenomena and provides insights into learners’ immediate emotional experiences. The instrument allows for easy customisation to target emotions specific to particular stimuli, appraisals, and action responses. While this methodology allows for capturing immediate, context-specific emotional reactions, if frequently used, it can become intrusive. Additionally, compared to techniques like image tracking or face-to-face interviews [77,78], this methodology is more reliable in assessing internal emotional states that may not be readily observable through external behaviours.

### 3.3 Associations between prior knowledge and emotions

Correlation analyses (Fig 4) revealed negative associations between prior knowledge level (pretest score) and specific negative anticipatory emotions of participants, including nervousness (Spearman correlation = −0.156; p = 0.031), frustration (Spearman correlation = −0.158; p = 0.029), and shame (Spearman correlation = −0.184; p = 0.011). This confirms our previous findings on the association between prior knowledge of microbiology and nervousness and frustration [58].

**Fig 4.**
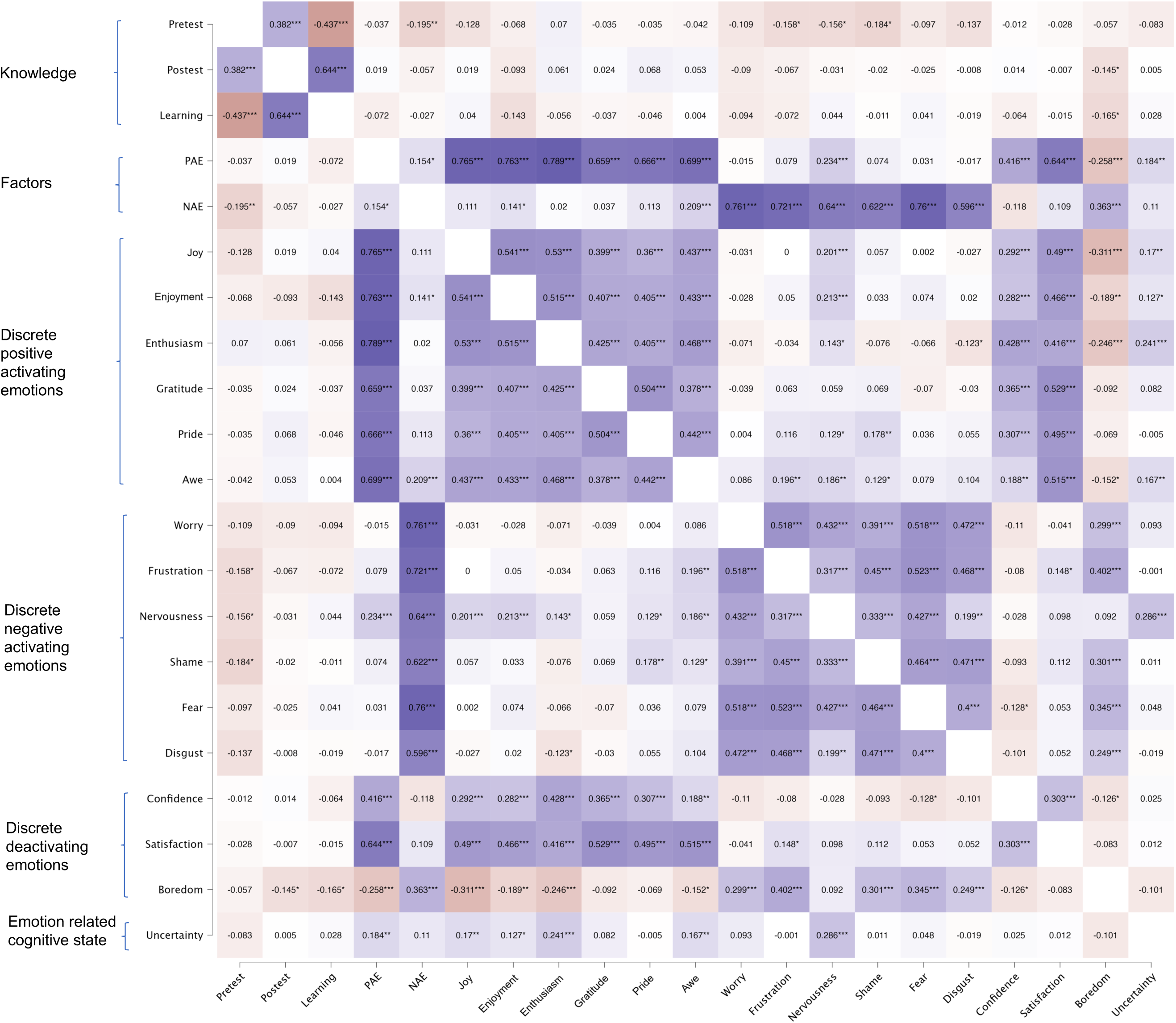
Heatmap of intercorrelations. *p<.05, **p<.01, ***p<.001

In agreement, correlation analyses of latent factors based on emotion valence (PAE or NAE) showed that the pretest score was negatively related to NAE (Spearman correlation = −0.195; p = 0.007), but not significantly associated with PAE (Fig 4). These correlations were not significantly different according to sex or prior education (Fig 5). Overall, these results indicated that students who anticipated greater aversion to the activity possessed less prior scientific knowledge. Since these anticipatory emotions were situational (a laboratory activity), and the scientific knowledge derived from prior education, we reasoned that anticipatory emotions were modulated either directly by prior education (as supported by Robust ANOVA) or mediated by the knowledge acquired at this educational stage (as supported correlations). A mediation analysis showed that the effect of prior education on NAE (prior education →NAE; Total effect β=0,258; p=0.027) was partially mediated by prior knowledge (prior education→prior knowledge →NAE; β=0,106 p=0.026). These direct and indirect effects of prior education on emotions were similar to those observed before a microbiology activity, but in this case, the effect was on PAE, not NAE [79]. These results indicated that preservice teachers with lesser scientific knowledge tend to anticipate more negative activating emotions (or less positive activating emotions) when facing the prospect of engaging in experimental science activities. Additionally, it unveils that an individual’s prior education significantly affects not only their level of prior knowledge but also their anticipatory emotions towards activities, in agreement with preceding studies that explored the impact of past academic performance on emotions and the known effect of perceived competence and control on momentary emotions [31,40,80,81]. Our results suggest that such perceptions of competence and control emerge not solely from students’ understanding of the subject matter but also from their awareness of the constraints imposed by their lack of a background in STEM. As we previously observed before a microbiology activity [79], anticipated students’ emotional experiences were influenced by their self-assessed knowledge and perceived limitations related to their educational background.

**Fig 5.**
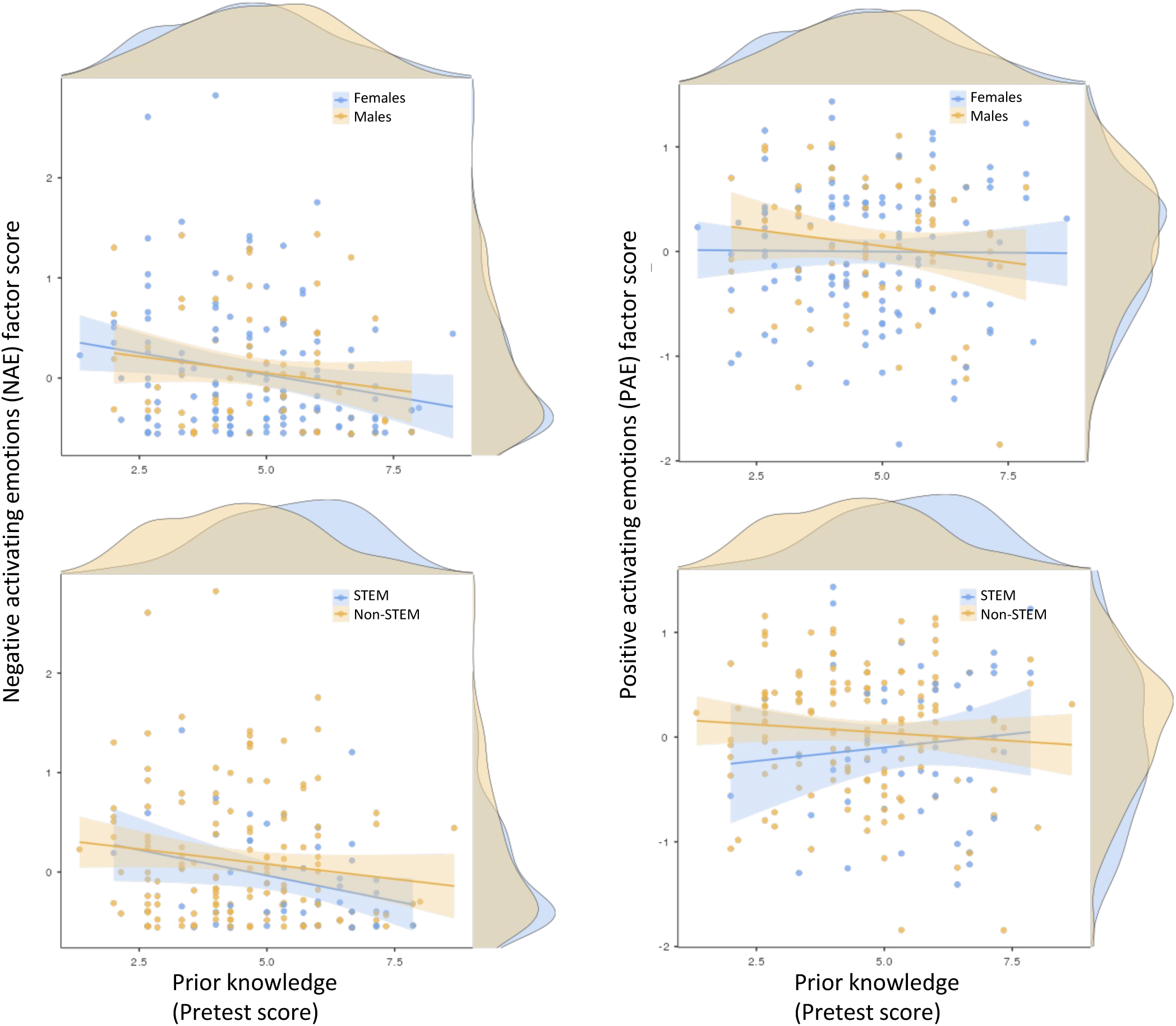
Linear regression analysis showing the relationship between pretest scores and the factor scores of anticipatory negative (left panels) and positive (right panels) activating emotions, segregated by sex (top panels) and prior education (bottom panels). The density map of data distribution is represented on the axis, while the shaded area on either side of the straight line represents the standard error of the mean.

Earlier research has demonstrated that low learning outcomes in areas such as languages and mathematics often precede negative emotions (including boredom, anger, shame, and despair) across various educational stages [23,37,38,81]. The current study supports the predictive nature of past performance concerning negative anticipatory emotions in preservice teachers. While investigations into the interplay between learning and emotion during specific (situational) science activities among preservice teachers remain limited, the underlying cause of negative emotions and low self-efficacy in science among trainee teachers within the Spanish educational milieu is hypothesised to stem from their insufficient science training and subsequent withdrawal from science curricula post-secondary education [2,82,83]. Previous research has also indicated that good academic results predict positive emotions such as enthusiasm, excitement, and pride [31,40,58,80,81]. However, this association was not observed in the present study.

### 3.4 Associations between anticipatory emotions, achievement, and learning

Prior knowledge correlates with achievement (Spearman correlation = 0.382; p <0.001) and learning (Spearman correlation = −0.437; p <0.001). According to linear regression analysis, prior knowledge explains 14.4% of the achievement variance [β=0.375±0.0674 (Avg±SE), p<0.001; R²=0.144], and explains 19.9% of learning variance [β=0.581±0.0946 (Avg±SE), p<0.001; R²=0.199]. The negative correlation between learning and prior knowledge indicates that participants with lower pretest scores benefit more from new learning, unlike those with higher scores who likely had this knowledge initially. Noteworthy, unlike prior knowledge and achievement variances, learning variances were not attributable to sex or prior education (Table 5). In our previous study of anticipatory emotions before a microbiology activity, joy and enthusiasm were positively associated with achievement, while frustration and concern were negatively associated [58]. In this study, the boredom of participants is the only discrete emotion associated with achievement (Spearman correlation = −0.145; p = 0.020) and learning (Spearman correlation = −0.165; p = 0.041). The ability of this anticipatory emotion to predict a decrease in achievement and learning was not unexpected since boredom reduces cognitive resources and task-related attention. The observed correlation coefficients for boredom’s impact on achievement are marginally below the threshold previously reported [32], which identified a stronger association within the range [−0.35, −0.25] in situational academic contexts. The mediation analysis revealed that boredom did not mediate the relationship between prior knowledge and learning outcomes or achievement. While previous studies have often reported this mediation effect, it appears to be non-uniform across different contexts. Recent evidence highlights this variability; for example,[84] observed that boredom’s mediation between cognitive appraisal and math achievement occurs in only about 47% of the educational systems examined. This underscores the complexity and situational nature of the mediating role of boredom in educational settings. Nevertheless, the analysis identified a significant mediating effect of boredom between negative anticipatory emotions (NAE) and educational outcomes: learning (NAE → boredom → learning; β = −0.118 ± 0.053 (Avg ± SEM), p = 0.025; R² = 0.255) and achievement (NAE → boredom → achievement; β = −0.076 ± 0.036 (Avg ± SEM), p = 0.037; R² = 0.219), indicating that boredom indeed plays a role in how negative anticipatory emotions affect both learning and achievement.

To deeply explore the relationship between achievement and boredom, we studied the relationship between prior knowledge and achievement at different boredom ratings (Fig 6). The histogram displays a clear inverse relationship between median boredom levels and academic achievement, with higher boredom correlating with decreased median achievement scores on the corresponding Likert scale. This trend is supported by linear regression analyses that consider both achievement and prior knowledge, revealing that boredom’s detrimental effect on achievement is accentuated among students with less prior knowledge. For instance, students with a high boredom rating of 4 and prior knowledge at 2.5 had an average achievement score of around 3. In contrast, those who did not anticipate boredom (rated 1 on the Likert scale) and identical prior knowledge tended to achieve significantly higher scores nearing 5. This pattern aligns with the correlation trend that higher achievement is associated with lower boredom.

**Fig 6.**
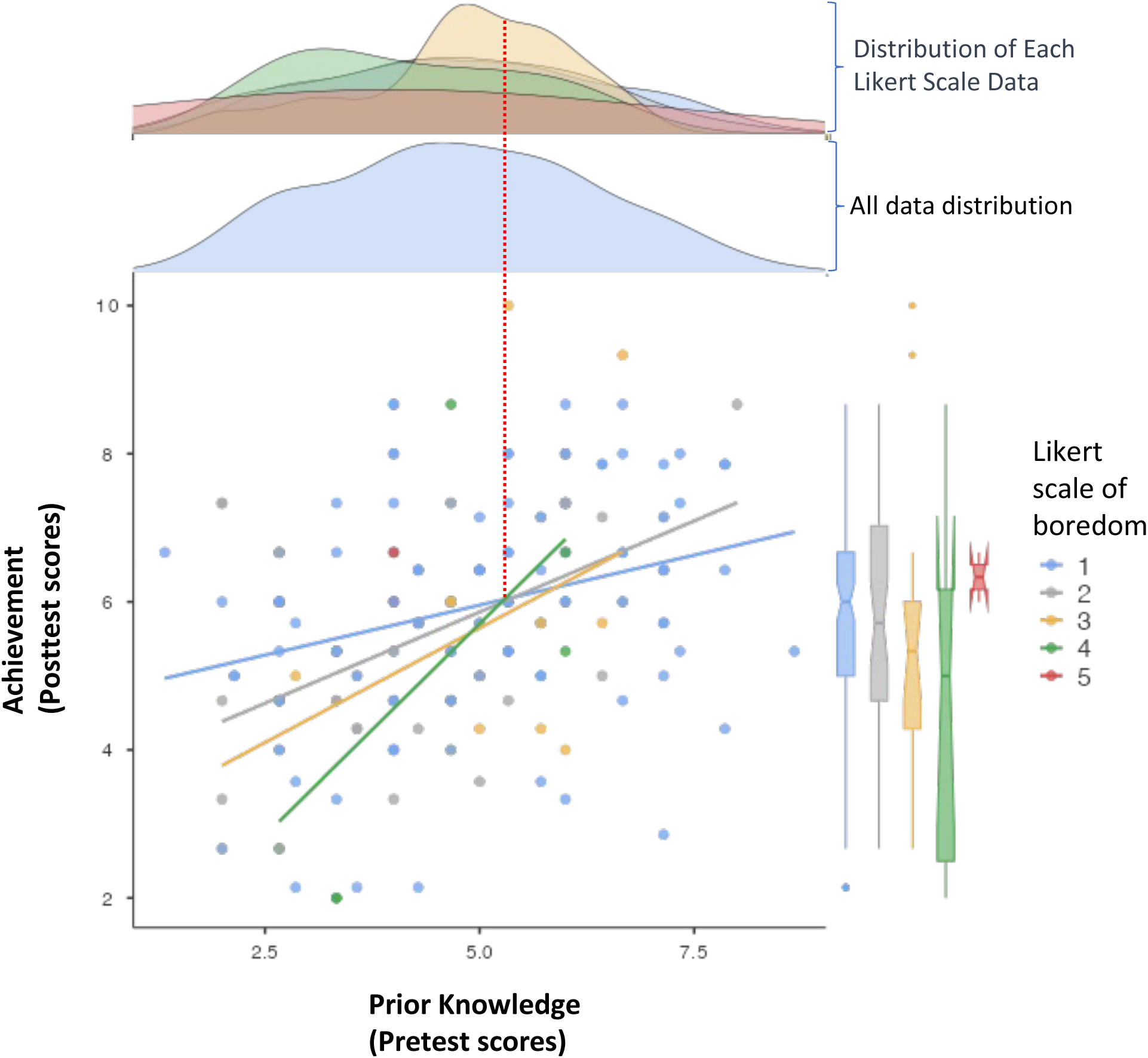
Scatter plot and linear regressions showing the impact of anticipated boredom levels on prior knowledge and achievement relationship. Boredom levels are rated on a Likert scale of 1 to 5 and depicted in varying colors of dots, lines, and boxes. Individual linear regression models represent the relationship between prior knowledge (pretest scores) and achievement (posttest scores) for each boredom level. The box plot shows the distribution of achievement across each boredom level, with the horizontal line within each box marking the median achievement (50th percentile) for that particular boredom level. Box-plot edges, 25th and 75th percentiles; whisker sizes, 1.5C×Cinterquartile range (IQR); dots under and over the boxes are single data points. The density maps above the core diagram illustrate the dispersion of prior knowledge data across the spectrum (blue map at the bottom) and within specific levels of boredom (top map). A red dashed line extends from the intercepts of the regression analyses to the density maps, drawing attention to the concentration of data points associated with lower levels of prior knowledge.

Nevertheless, the achievement gap between different levels of boredom narrows with increasing prior knowledge, which diminishes the strength of the negative relationship between achievement and boredom. Notably, an inflexion point is observed in the data where boredom’s impact on achievement reverses; beyond a certain level of prior knowledge (denoted by the intercept of regression lines in Fig 6), the relationship becomes positive—higher boredom corresponds with higher achievement. This counterintuitive effect, the reversal of boredom’s impact, may explain the variability in the boredom-achievement correlation reported in the literature [32], as it hinges on the level of prior knowledge. The prevalence of a negative correlation between boredom and achievement in our analysis can be attributed to the higher representation of participants with lower prior knowledge within our sample (separated by a red dashed line in the density maps of Fig 6). The differential effect of boredom fully agrees with boredom’s mediating and moderating role in multiple-text comprehension [34], and we describe it for the first time in a science education context.

To methodically examine this relationship, we conducted a moderation analysis to discern how boredom anticipation influences the nexus between achievement and prior knowledge (Fig 7). The analysis confirms that prior knowledge is a significant predictor of achievement [β=0.381±0.0698 (Avg±SE), p<0.001], with the moderating effect of boredom substantially reducing this influence to β=0.2090±0.0950 (Avg±SE), (p=0.022). Further scrutiny of the moderation effect across various discrete emotions and states revealed that shame [β=0.1990±0.00817 (Avg±SE), p=0.015], and uncertainty [β=0.1466±0.0561 (Avg±SE), p=0.009], have a significant moderating impact on the relationship between pretest and posttest scores (Table 8). This comprehensive approach reinforces the importance of prior knowledge in academic achievement and highlights the multifaceted role that emotional states play in this dynamic.

**Fig 7.**
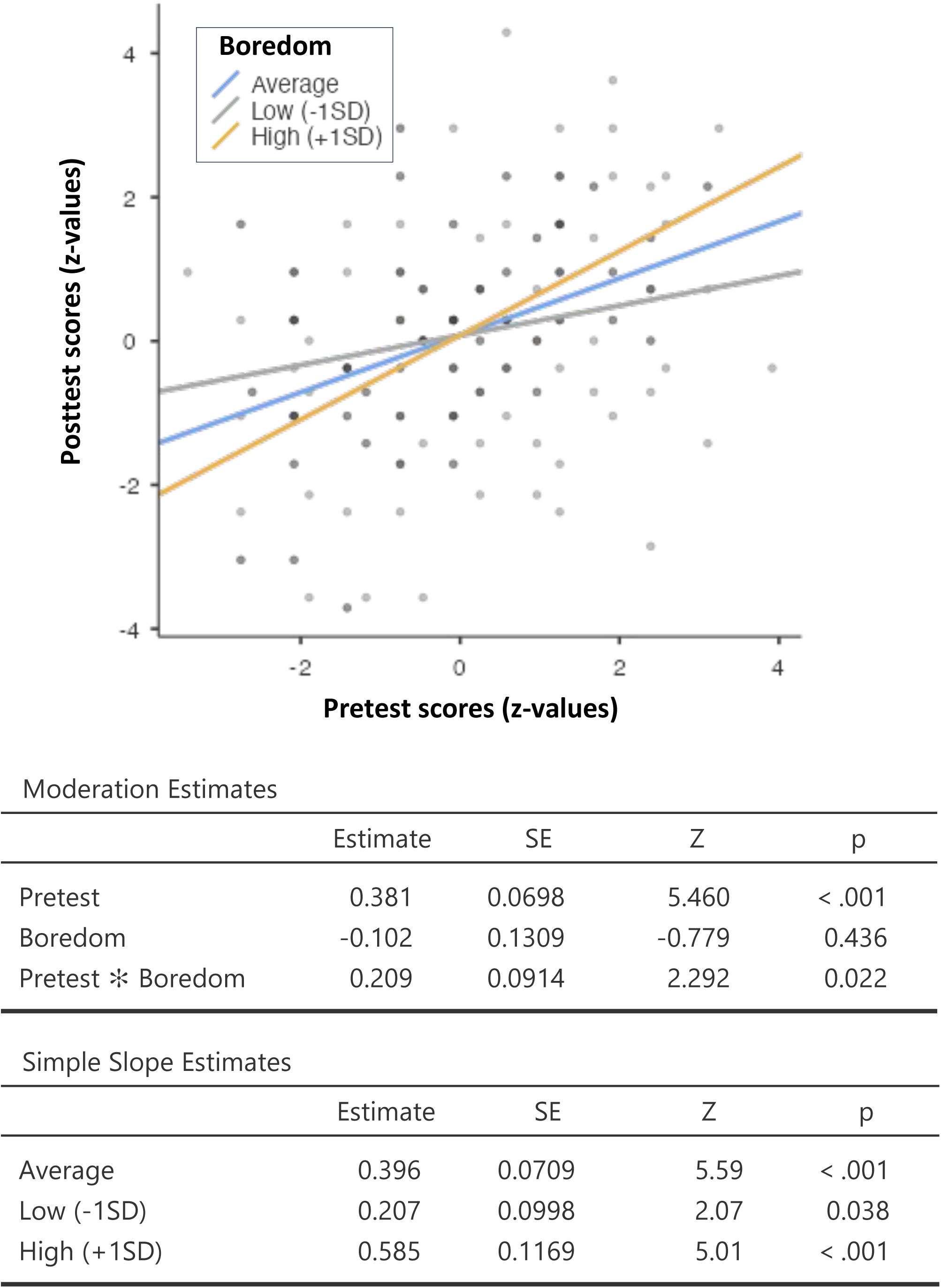
Moderation analysis of boredom’s influence on the relationship between Pretest (prior knowledge) and Posttest (achievement). The graph depicts the linear regression predicting Posttest scores based on Pretest scores at varying levels of the Boredom moderator (Average, Low (−1SD), and High (+1SD). The moderation estimates table (top) presents the effect of pretest scores on posttest, independent of boredom; the direct influence of boredom on posttest scores, without considering pretest scores; and the interaction effect (Pretest * Boredom), illustrating how the Pretest-Posttest relationship varies with different boredom levels. The simple slope estimates table (bottom) details the relationship between Pretest and Posttest scores for specific boredom levels.

**Table 8.**
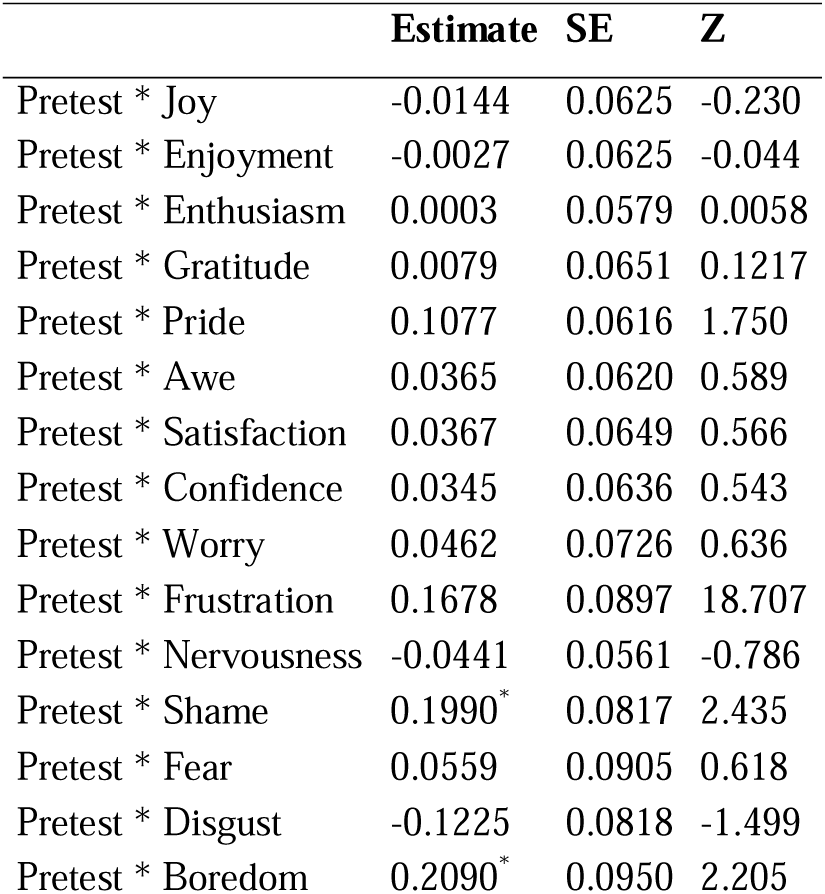
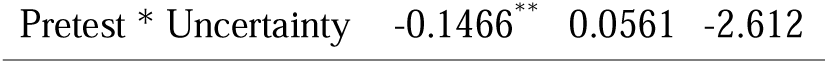
Moderation analysis of discrete feelings on the relationship between prior knowledge and achievement. (* p-value≤0.05, ** p-value≤0.01).

These observations are consistent with previous studies highlighting the predictive value of anticipatory emotions on students’ academic performance across various educational stages. In particular, the negative influence of boredom aligns with studies demonstrating that high prior intensity of this negative emotion predicts poor learning outcomes [5,30–32,40,85]. Previous research has also identified the predictive value of other positive and negative emotions [57,58], although these interactions were not observed in the present study.

This study extends prior research by exploring the role of anticipatory emotions, particularly boredom, in preservice teachers’ science learning experiences. Our findings align with established theories, such as the Control-Value Theory, demonstrating the significant impact of emotions on learning outcomes [1]. In particular, our work underscores the detrimental effect of boredom on academic achievement [5,17,21,22], especially for students with lower prior knowledge [34]. Addressing this emotion early in the educational process can potentially improve subsequent learning outcomes [38]. Interestingly, boredom does not directly mediate the effect of prior knowledge on achievement but rather operates through NAE, which is associated with non-STEM backgrounds and low prior knowledge. Low prior knowledge increases NAE, which in turn increases boredom, exerting a negative effect on achievement. These results suggest that interventions should not only aim to counteract boredom but also address the lack of confidence and science knowledge of non-STEM students, which are required to perform the activity effectively.

### 3.5 Interplay of emotions and learning

Statistical network analysis [71] integrated the relationships between anticipatory emotions and prior knowledge and learning, providing a systemic perspective of the relationships between these variables (Fig 8). The network’s sparsity coefficient, at 0.627, indicates a high level of unconnectedness among potential variable pairings, except for three cluster structures. The first cluster encompasses NAE and boredom (sparsity=0.19), suggesting closer interaction within this grouping. The second cluster comprises PAE, satisfaction and confidence (all positive emotions) with dense interconnectivity (sparsity=0.143). The proximity of PAE and NAE, discrete emotions in each cluster, is consistent and adds validity to the two-factor model for PAE and NAE scales. The third, more diminutive cluster, consists of cognitive elements (prior knowledge and learning outcomes). The relatively low connectivity between PAE and NAE clusters agrees with the low correlation of these scales (Spearman correlation =0.154, p=0.011). The observed network structure aligns with Pekrun’s taxonomy of emotions [17], where affective states are grouped according to their valence and activation properties. Notably, nervousness, though proximal to the negative emotions cluster, occupies a peripheral position, likely attributable to its classification as an emotional state, as opposed to a discrete emotion like fear, shame, or disgust. The distant positions of boredom and confidence from NAE and PAE, respectively, corroborate its characterisation as deactivating emotions. However, the central placement of satisfaction within PAE cluster presents an anomaly, given its classification as a positive deactivating emotion [17]. This suggests that the anticipation of satisfaction has an activating character.

**Figure 8.**
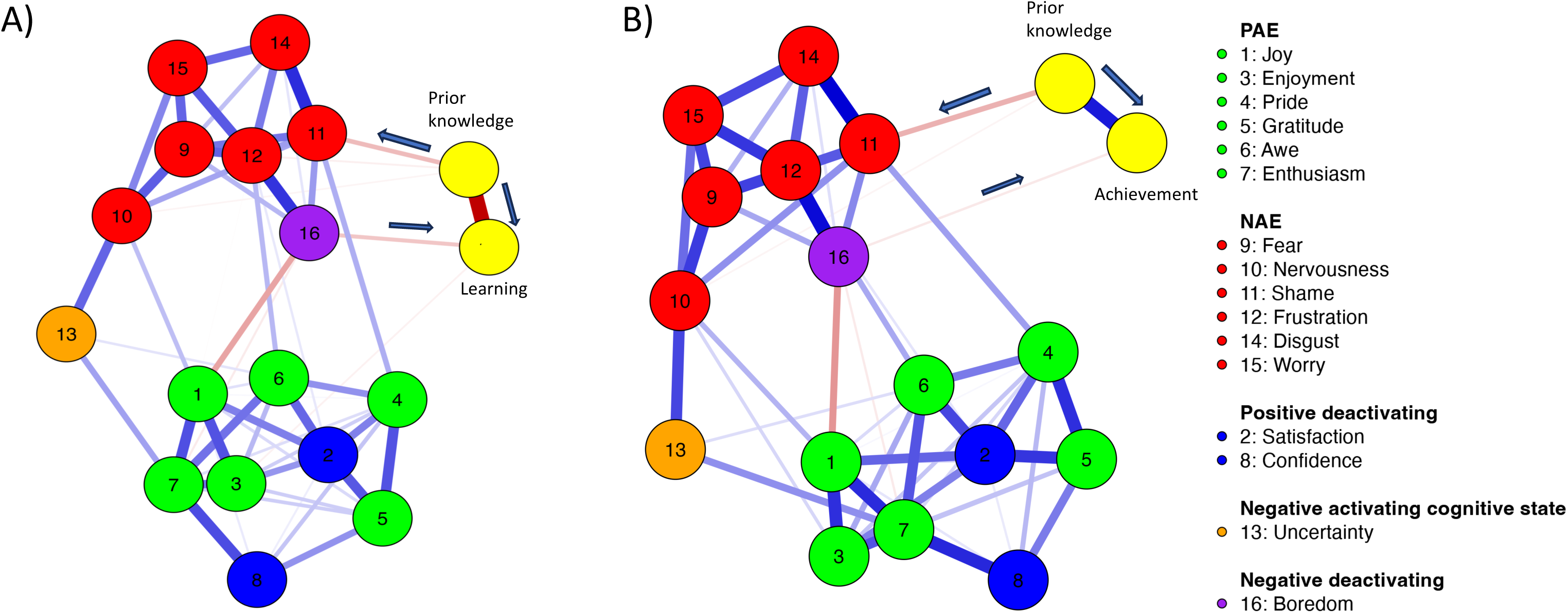
Network plots of variables considering A) prior knowledge and learning and B) prior knowledge and achievement. Nodes (circles) represent variables identified by numbers in the left margin, except prior knowledge, achievement and learning, shown close to their yellow nodes. Edge weights (lines) represent the strength of association between variables. Blue edges represent positive associations. Red edges represent negative associations. Denser lines represent stronger connections. Arrows indicate the direction of interactions according to this study.

The network also captures expected intra-cluster positive associations within positive and negative emotions and the anticipated negative interplay between boredom and joy. Intriguing is the positive correlation between enthusiasm and uncertainty, which may be attributable to the novelty and excitement of participants engaged for the first time in practical DNA extraction using household materials. The positive association between awe and uncertainty also supports this possibility. Additionally, the positive link between pride and shame, emotions with cognitive antecedents of success or failure, respectively, and traditionally seen as opposite [17], suggests that participants may anticipate the feeling of shame they believe they can overcome. The network mappings corroborate relationships identified in our previous study through alternative analytical approaches within this research, specifically the impacts of shame, frustration, and nervousness on prior learning and the influence of boredom on both achievement and learning [79]. Furthermore, the networks reveal an unforeseen negative link between enjoyment and academic achievement. This association could reflect the potentially disruptive role of excessive enjoyment in learning contexts, possibly impeding achievement by diverting attention from task-related goals [18], which is consistent with the elevated excitement levels discussed earlier.

The results from the network analysis indicate that while negative emotions exhibit mutual interactions and modulation effects, it is predominantly boredom that mediates their impact on learning and achievement. Likewise, while positive emotions also show interconnectivity and modulation effects among themselves, enjoyment emerges as the principal emotion influencing learning. These insights present a notable divergence from the emotion-learning interplay before a microbiology laboratory session [58]. In that study, despite the overall similarity of the network (the observed interplay and modulatory influence among both negative and positive emotions, and the similar effect of frustration and nervousness on prior knowledge), it was specifically frustration that negatively affected achievement, while nervousness had a positive impact. These findings underscore the relevance of contextual investigation into emotional experiences to forecast emotional responses more accurately and potentially strategically influence them in an educational setting.

## 4. Concluding remarks

This study serves as a conceptual replication of prior research, examining whether the well-established impact of emotions on learning outcomes—traditionally documented in the context of courses and grades—is generalisable to preservice teachers engaged in laboratory activities. Results align with several previous research findings, including (i) the strong intercorrelation among negative emotions and within positive emotions (Fig 4) [22,50,57]; (ii) the latent factors behind negative and positive activating emotions [58,86]; (iii) the minimal correlation between these factors [22,48,57];(iv) the measurement invariance across sex and prior education [40]; (v) the inverse relationship between negative emotions and both prior knowledge and achievement [48,87,88]; and (vi) the mediating role of boredom in achievement and learning [31,32,34]. This study supports the extension of prior research to preservice teachers’ education and adds validity to our emotions assessment tool.

Our framework for categorising emotional reactivity components offers precise descriptions of focus elements, valuation, and actions (Fig. 1), facilitating comparability across studies. By identifying the triggers of emotions in specific classroom activities (Table 2), we enhance the potential for comparison and generalisation, particularly in course-specific contexts with numerous variables (Fig. 2). This categorisation moves beyond mere associations between emotions and academic achievement, paving the way for uncovering causal relationships and enabling the systematic study of emotional phenomena.

We also present an experience sampling method that is minimally disruptive, allowing for quick emotional assessments during classroom activities (Fig. 2). Our replication study supports the reliability and validity of this instrument, offering valuable insights for teachers and researchers to adapt teaching practices based on emotional data. Notably, emotions like gratitude, shame, disgust, worry, and uncertainty vary by sex and prior education, consistent with research focused on specific situational contexts [44,89,90] and overall course performance [39]. This supports the generalisation of findings across different time frames. Moreover, the single-item instrument to assess discrete emotions offers flexibility, enabling practitioners to customise it to diverse educational contexts. For instance, disgust, often overlooked, was included in this study due to its potential elicitation during DNA extraction practices.

Our findings further corroborate that emotions are influenced by prior academic experiences. Students with higher prior knowledge anticipated fewer negative emotions, suggesting that prior knowledge directly influences emotional responses during educational activities. Addressing these emotional patterns can guide the design of effective emotional regulation strategies, fostering a conducive learning environment. Since emotions can be regulated over time [37], targeted interventions are possible.

In line with recent research [23], boredom in specific classroom tasks has an immediate effect on academic performance. However, addressing boredom before the activity begins can mitigate its detrimental impact. Since boredom often arises from low perceived control or low value placed on the task [34,35], highlighting the utility and accessibility of the task—even for inexperienced students—could help neutralise the negative effects of boredom, thereby improving engagement and performance.

A key contribution of our research is integrating network analysis with traditional methods such as correlation, latent factor, mediation, and modulation analyses. Our network analysis elucidates the interrelationships between variables, mapping them based on their shared covariance. Within the network, boredom emerges as a pivotal element, influenced by NAE, inhibited by PAE, and negatively impacting learning and achievement. This central positioning of boredom aligns with findings from mediation and modulation analyses. Antecedents like prior education and learning appraisals, which negatively predict NAE, underscore the potential of interventions aimed at reshaping these appraisals—especially for non-STEM students—to enhance achievement. Furthermore, our results advocate for strategies that mitigate negative emotions, such as modifying experimental materials to decrease disgust, to foster better learning outcomes. Given that emotions are amenable to intervention, our findings propose that specifically addressing boredom in students with low prior knowledge could amplify the benefits of educational strategies.

In our prior research on emotions preceding a microbiology activity [58], boredom was similarly situated centrally between NAE and PAE. Yet, in that context, it was frustration, rather than boredom, that detrimentally impacted learning. Moreover, in that study, nervousness (which used to negatively impact learning outcomes) had a counterintuitive positive effect on achievement. Here, we did not observe this association, but we observed another counterintuitive interaction, a negative association between enjoyment and learning. These non-univalent interactions of emotions and learning have been previously described [18] and underscore the importance of investigating the specific emotional dynamics and their regulatory mechanisms across various classroom activities to tailor emotional regulation strategies more effectively. It is essential to recognise that the impact of emotions on learning is not solely dependent on their valence. Some positive emotions, like enjoyment, can be detrimental to learning if experienced with high intensity or unrelated to specific scientific content. Likewise, a certain intensity of some stimulating negative emotions (e.g., nervousness) can enhance learning. During initial training, these factors should be considered.

Moreover, results support existing psychological and neuroscientific theories on the bidirectional emotion-cognition interaction and demonstrate its presence in a sample of future teachers in science education. The detected interactions highlight the importance of prior science learning experiences in shaping the anticipatory emotions experienced by preservice teachers, which influence their science learning during initial training. In the context of constructed emotion theory, emotional responses are predictive, drawing from past experiences in analogous situations to inform future behaviour. Anticipatory emotions thus act as a prism through which we can view a student’s previous experiences, whether favourable or adverse, and gauge the corresponding level of arousal, which encompasses attention, motivation, and memory. This predictive quality of emotional responses is crucial for educational practitioners, as it facilitates tailoring classroom strategies from the outset. By reviewing and reinforcing requisite prior knowledge at the class’s beginning, educators can ensure the accessibility of learning activities, regardless of students’ educational backgrounds. The application of this straightforward pedagogical strategy could be instrumental in altering students’ anticipatory emotional states, thereby reshaping their predictive landscape and potentially enhancing educational outcomes.

In conclusion, the present study reveals the complex interplay of anticipatory emotions, prior knowledge, and learning outcomes in preservice teachers. It underscores the importance of considering both positive and negative emotions in the teaching-learning process and the need for teachers to effectively manage and regulate their students’ emotions. By recognising the potential effects of emotions on learning, teachers can adapt their teaching methods and classroom environments to foster an atmosphere that supports learning and emotional well-being.

## 5. Limitations and future research directions

This study has several limitations. First, the participants were from the same context (same region, similar educational backgrounds, and the same training program), limiting the generalizability of the results. Second, the study relied on a quantitative self-report test of emotions, which measures feelings rather than actual emotions and might be subject to individual interpretation. Future research could use open-ended questionnaires or interviews for qualitative analysis to further explore the relationship between anticipatory emotions, prior knowledge, and other aspects of learning. The work is based on the assumption that participant expresses their emotions in relation to a focus (activity), but their expression can be related to other focuses.

Future research should further explore the relationships between different emotions and learning outcomes and investigate teachers’ strategies to manage and regulate their students’ emotions effectively. Additionally, longitudinal studies can help clarify anticipatory emotions’ long-term effects on learning and academic achievement in various educational contexts.

## Acknowledgments

This study has been supported by Grant PID2022-139684NB-I00 funded by MCIN/AEI/ 10.13039/501100011033 and by "ERDF A way of making Europe".

## Author contributions

Conceptualisation: JAGOA, JMMM, and MREG

Formal analysis: JAGOA, JMMM, CVM, and MREG

Supervision: JAGOA, JMMM, and MREG

Methodology: JAGOA, JMMM, and MREG

Writing – original draft: JAGOA and JMMM

Writing – review & editing: JAGOA, JMMM, CVM, and MREG

## Declaration of generative AI and AI-assisted technologies in the writing process

During the preparation of this work the authors used with caution ChatGPT4 and Grammarly to improve language and readability. After using these services, the authors reviewed and edited the content as needed and take full responsibility for the content of the publication.

## Supporting information

S1 File. Dataset.csv

Dataset containing the information collected from participants and utilised in the analysis.

