## Supplementary material for "Anticipatory emotions and academic performance: The role of boredom in a preservice teachers’ lab experience"

Instrument original Version (Spanish) 2

Instrument English translation 5

References 7

**Instrument original Version (Spanish)**

**Práctica de extracción de ADN con material cotidiano**
Has sido invitado/a participar en un estudio de investigación dirigido a mejorar la enseñanza en la Universidad de Extremadura. El estudio trata de comprender la relación entre el aprendizaje y la percepción que tiene el alumno de las actividades que se desarrollan en el aula. En este caso, la extracción de ADN con material cotidiano. En las futuras publicaciones los datos obtenidos se mostrarán globalmente, preservando completamente las características individuales de los participantes, que serán siempre anónimas. La participación es voluntaria, no tiene ningún riesgo y no afecta de ninguna manera la calificación de los participantes. El proyecto dura 3 años y no tiene ánimo de lucro. El/la participante puede abandonar en cualquier momento.

He leído y
- ○ Acepto participar
- ○ Prefiero no continuar

**Datos personales**
Escribe una clave formada por: inicial del nombre de tu madre, inicial del nombre de tu padre, dos últimas cifras y letra de tu DNI (por ejemplo: MJ51A) __________________

¿Cómo te identificas?
- ○ Mujer
- ○ Hombre
- ○ Otros

¿Cómo accediste al Grado de Maestro en Educación Primaria?
- ○ Bachillerato de Ciencias
- ○ Bachillerato de Humanidades o Ciencias Sociales
- ○ Bachillerato de Arte
- ○ Formación Profesional
- ○ Otra

**Parte 1. Las emociones hacia una actividad práctica** (Marcos-Merino *et al.* 2020)

¿Qué emociones anticipas sentir al enfrentarte a una práctica de extracción de ADN utilizando materiales cotidianos?

Señala la intensidad desde "1 "nada" a 5 "mucho o intensamente"

| | 1 | 2 | 3 | 4 | 5 |
|---|---|---|---|---|---|
| Alegría | ○ | ○ | ○ | ○ | ○ |
| Preocupación | ○ | ○ | ○ | ○ | ○ |
| Confianza | ○ | ○ | ○ | ○ | ○ |
| Frustración | ○ | ○ | ○ | ○ | ○ |
| Satisfacción | ○ | ○ | ○ | ○ | ○ |
| Incertidumbre | ○ | ○ | ○ | ○ | ○ |
| Entusiasmo | ○ | ○ | ○ | ○ | ○ |
| Nerviosismo | ○ | ○ | ○ | ○ | ○ |
| Diversión | ○ | ○ | ○ | ○ | ○ |
| Aburrimiento | ○ | ○ | ○ | ○ | ○ |
| Gratitud | ○ | ○ | ○ | ○ | ○ |
| Asco | ○ | ○ | ○ | ○ | ○ |
| Orgullo | ○ | ○ | ○ | ○ | ○ |
| Vergüenza | ○ | ○ | ○ | ○ | ○ |
| Sorpresa | ○ | ○ | ○ | ○ | ○ |
| Miedo | ○ | ○ | ○ | ○ | ○ |

**Parte 2. Responde a las siguientes preguntas, solo una opción correcta** (Ochoa de Alda *et al.* 2019)

1. Podemos extraer material genético de:
a) Solo los seres vivos
b) Solo los procariotas
c) Solo los eucariotas
d) De todos los seres vivos y algunos no vivos (por ejemplo, virus)

2. ¿Dónde crees que se encuentra el ADN en un tomate?:
a) Solo en las células de la semilla
b) Solo en las células de la pulpa (del fruto)
c) Solo en las células de la piel
d) En todas las células

3. El ADN es:
a) Un ácido nucleico
b) Una grasa rodeada de fósforo
c) Una proteína
d) Un aminoácido

4. En los humanos los cromosomas sexuales se encuentran exclusivamente en:
a) Las células de los testículos y los ovarios
b) Todas las células con núcleo
c) Los espermatozoides y los óvulos
d) Las mitocondrias de todas las células

5. Qué tipos de células tienen mitocondrias y cloroplastos:
a) Las células animales
b) Las células vegetales
c) Las bacterias
d) Ninguna, o bien tienen mitocondrias o bien tienen cloroplastos

6. La diferencia esencial entre célula procariota y célula eucariota radica en:
a) El tamaño celular
b) La pared celular
c) El núcleo celular
d) La composición química del citoplasma

7. Respecto al ADN señala la respuesta CORRECTA:
a) Se encuentra exclusivamente en el núcleo celular
b) Es el único ácido nucleico de la célula
c) Se encuentra en cloroplastos, mitocondrias y núcleo
d) Está formado por aminoácidos

8. Señala la afirmación CORRECTA:
a) Todas las células están rodeadas por una membrana plasmática rígida
b) La membrana plasmática está formada fundamentalmente por lípidos, proteínas, carbohidratos y nucleótidos
c) Dentro de las células algunos orgánulos están rodeados por membranas similares a la membrana plasmática
d) Solo las células eucariotas tienen membrana plasmática

9. En los vegetales, la pared celular es:
a) Una estructura extracelular formada por polisacáridos, fundamentalmente celulosa
b) Una estructura rígida que las rodeas
c) La materia prima para formar el papel
d) Todas las respuestas son correctas

10. Identifica la relación «tipo celular - características» INCORRECTA:
a) Todas las células vegetales tienen pared celular
b) Todas las células animales tienen mitocondrias y cloroplastos
c) Todas las células vegetales tienen mitocondrias y cloroplastos
d) Todas las células animales tienen mitocondrias

11. Los riñones son órganos que se encuentran en el cuerpo humano. A un hombre le sacaron uno de sus dos riñones cuando era joven porque estaba enfermo. Ahora tiene un hijo. ¿Cuántos riñones tuvo su hijo al nacer?
a) 1, ya que solo se puede heredar lo que se tiene
b) 2, ya que no se han alterado sus genes
c) 2, ya que los riñones se regeneran
d) 1 o 2 según si hereda 1 o ninguno del padre y el de la madre

12. ¿Cuál de las siguientes es la mejor descripción del propósito de la respiración celular?
a) Proporcionar energía para la actividad celular
b) Producir azúcar para almacenar en las células
c) Liberar oxígeno para la respiración
d) Proporcionar dióxido de carbono para la fotosíntesis

13. ¿Qué tipo de células destruyen a las bacterias que invaden el cuerpo?
a) Los glóbulos blancos
b) Los glóbulos rojos
c) Las células del riñón
d) Las células del pulmón

14. La imagen muestra una célula. ¿Cuál es la función de la parte de la célula marcada con una X?
a) Almacenar agua
b) Producir alimento
c) Absorber energía
d) Controlar las actividades

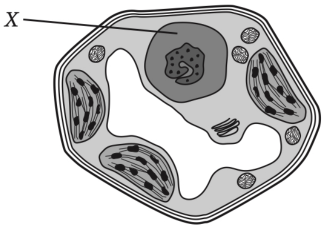

**Instrument English translation**

**Perception of a DNA Extraction Practice with Everyday Materials**
You have been invited to participate in a research study to improve teaching at the University. The study seeks to understand the relationship between learning and the student's perception of the activities carried out in the classroom. In this case, DNA extraction with everyday materials. In future publications, the data obtained will be presented globally, fully preserving the individual characteristics of the participants, who will always remain anonymous. Participation is voluntary, poses no risk, and does not affect the participants' grades in any way. The project lasts for three years and is not for profit. The participant may withdraw at any time.

I have read and
- ○ I agree to participate.
- ○ I prefer not to continue.

**Personal Information**
Write a code formed by the first initial of your mother's name, the first initial of your father's name, the last two digits and the letter of your ID (for example MJ51A) ________________

How do you identify yourself?
- ○ Female
- ○ Male
- ○ Other

How did you access the Bachelor's Degree in Primary Education?
- ○ Science Track High School
- ○ Humanities or Social Sciences Track High School
- ○ Art Track High School
- ○ Vocational Training
- ○ Other

**Part 1. Emotions towards a practical activity** (Marcos-Merino *et al.* 2020)

What emotions do you anticipate feeling when confronted with a DNA extraction practice using everyday materials?

Indicate the intensity from "1" none at all to "5" a lot or intensely

| | 1 | 2 | 3 | 4 | 5 |
|---|---|---|---|---|---|
| Joy | ○ | ○ | ○ | ○ | ○ |
| Worry | ○ | ○ | ○ | ○ | ○ |
| Confidence | ○ | ○ | ○ | ○ | ○ |
| Frustration | ○ | ○ | ○ | ○ | ○ |
| Satisfaction | ○ | ○ | ○ | ○ | ○ |
| Uncertainty | ○ | ○ | ○ | ○ | ○ |
| Enthusiasm | ○ | ○ | ○ | ○ | ○ |
| Nervousness | ○ | ○ | ○ | ○ | ○ |
| Fun | ○ | ○ | ○ | ○ | ○ |
| Boredom | ○ | ○ | ○ | ○ | ○ |
| Gratitude | ○ | ○ | ○ | ○ | ○ |
| Disgust | ○ | ○ | ○ | ○ | ○ |
| Pride | ○ | ○ | ○ | ○ | ○ |
| Shame | ○ | ○ | ○ | ○ | ○ |
| Awe | ○ | ○ | ○ | ○ | ○ |
| Fear | ○ | ○ | ○ | ○ | ○ |

**Part 2. Answer the following questions, only one correct option** (Ochoa de Alda *et al.* 2019)

1. We can extract genetic material from:
a) Only living beings
b) Only prokaryotes
c) Only eukaryotes
d) From all living beings and some non-living (for example, viruses)

2. Where do you think DNA is located in a tomato?:
a) Only in the seed cells
b) Only in the pulp cells (of the fruit)
c) Only in the skin cells
d) In all cells

3. DNA is:
a) A nucleic acid
b) A fat surrounded by phosphorus
c) A protein
d) An amino acid

4. In humans, the sex chromosomes are found exclusively in:
a) The cells of the testicles and ovaries
b) All cells with a nucleus
c) The sperm and egg cells
d) The mitochondria of all cells

5. What types of cells have mitochondria and chloroplasts:
a) Animal cells
b) Plant cells
c) Bacteria
d) None. They either have mitochondria or chloroplasts

6. The essential difference between prokaryotic and eukaryotic cells lies in:
a) Cell size
b) The cell wall
c) The cell nucleus
d) The chemical composition of the cytoplasm

7. Regarding DNA, indicate the CORRECT answer:
a) It is found exclusively in the cell nucleus
b) It is the only nucleic acid in the cell
c) It is found in chloroplasts, mitochondria, and the nucleus
d) It is made up of amino acids

8. Indicate the CORRECT statement:
a) A rigid plasma membrane surrounds all cells
b) The plasma membrane is fundamentally composed of lipids, proteins, carbohydrates, and nucleotides
c) Within cells, some organelles are surrounded by membranes similar to the plasma membrane
d) Only eukaryotic cells have a plasma membrane

9. In plants, the cell wall is:
a) An extracellular structure formed mainly by polysaccharides, especially cellulose
b) A rigid structure that surrounds them
c) The raw material to make paper
d) All answers are correct

10. Identify the INCORRECT "cell type - characteristics" relationship:
a) All plant cells have a cell wall
b) All animal cells have mitochondria and chloroplasts

c) All plant cells have mitochondria and chloroplasts
d) All animal cells have mitochondria

11. The kidneys are organs found in the human body. A man had one of his two kidneys removed when he was young because it was sick. Now he has a son. How many kidneys did his son have at birth?
a) 1, since you can only inherit what you have
b) 2, since his genes have not been altered
c) 2, since kidneys regenerate
d) 1 or 2 depending on whether he inherits 1 or none from the father and one from the mother

13. ¿Qué tipo de células destruyen a las bacterias que invaden el cuerpo?
a) Los glóbulos blancos
b) Los glóbulos rojos
c) Las células del riñón
d) Las células del pulmón

14. La imagen muestra una célula. ¿Cuál es la función de la parte de la célula marcada con una X?
a) Almacenar agua
b) Producir alimento
c) Absorber energía
d) Controlar las actividades

12. Which of the following is the best description of the purpose of cellular respiration?
a) To provide energy for cellular activity
b) To produce sugar to store in cells
c) To release oxygen for breathing
d) To provide carbon dioxide for photosynthesis

13. What type of cells destroy bacteria invading the body?
a) White blood cells
b) Red blood cells
c) Kidney cells
d) Lung cells

14. The image shows a cell. What is the function of the part of the cell marked with an X?
a) Store water
b) Produce food
c) Absorb energy
d) Control activities

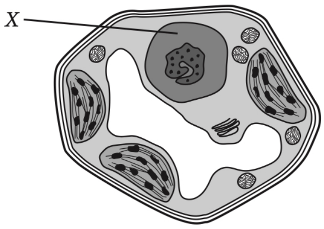
